# Membrane Phase, Charge, and Curvature Regulate α-Synuclein Binding Dynamics

**DOI:** 10.64898/2026.05.12.724662

**Authors:** Orianna H. Kou, Cailyn M. Sakurai, Stephanie Y. Ramirez, Brian H. Kim, David H. Johnson, Zixin Zhang, Christopher T. Lee, Wade F. Zeno

## Abstract

α-Synuclein (αSyn) is an intrinsically disordered protein whose interactions with lipid membranes are central to both its physiological function and its role in synucleopathies. While membrane charge, phase, and curvature are each known to influence αSyn binding, these properties are typically examined independently, leaving their combined effects on both equilibrium and dynamic membrane association unresolved. Here, we systematically investigate how membrane phase and charge jointly regulate αSyn binding, curvature sensitivity, and exchange dynamics using fluorescence microscopy, circular dichroism spectroscopy, and fluorescence recovery after photobleaching (FRAP), complemented by coarse-grained molecular dynamics simulations. Under zwitterionic conditions, αSyn preferentially binds highly curved gel-phase membranes, driven by curvature-dependent enrichment of packing defects arising from faceted vesicle morphologies. Incorporation of anionic lipids selectively enhances binding in liquid-phase membranes while attenuating curvature-dependent partitioning in gel-phase membranes. Dynamic measurements reveal that membrane phase and charge also govern the stability of membrane-associated αSyn, with gel-phase membranes and anionic lipids promoting kinetically stabilized states. Simulations show that curvature-induced defect formation is strongly amplified in gel-phase membranes but largely insensitive to charge. These findings establish that αSyn-membrane interactions are governed by a cooperative interplay between membrane phase, curvature, and charge and highlight the importance of resolving both thermodynamic and kinetic contributions to protein-membrane binding.

## Introduction

α-Synuclein (αSyn) is a small intrinsically disordered protein localized to the synaptic terminals of neurons^1^ and is closely associated with synucleopathies such as Parkinson’s Disease and Lewy body dementia^2–5^. Although its physiological function is not fully resolved, αSyn is known to interact with synaptic vesicle membranes, mitochondrial membranes, and plasma membranes^6–10^, where it has been implicated in vesicle trafficking and membrane remodeling^7;9;11–16^. These interactions are central to both its normal function and its pathological aggregation, motivating a detailed understanding of the physicochemical determinants that govern αSyn-membrane association.

Extensive work has established that αSyn undergoes a disordered-to-helix transition upon membrane binding, forming an amphipathic alpha helix within its N-terminal domain (NTD) that associates with lipid bilayers while the C-terminal domain (CTD) remains disordered and tethered to the membrane^17;18^. Electrostatic interactions between the net positively charged NTD and anionic lipid headgroups are well known to enhance binding^17;19–23^. In parallel, prior studies have reported enhanced αSyn association with gel-phase membranes relative to liquid-phase membranes^24–29^. This behavior is of particular relevance in the context of aging, as cellular membranes become more ordered due to increased lipid saturation and reduced polyunsaturation^30–32^. αSyn is also sensitive to membrane curvature, with preferential association to highly curved membranes such as small vesicles^19;26;28;33; 34^.

While membrane charge, phase, and curvature have each been shown to influence αSyn binding, these factors are typically considered independently. Few, if any, studies have systematically compared these effects simultaneously. As a result, it remains unclear how membrane phase and charge jointly regulate αSyn binding across curvature regimes. Moreover, the relative contributions of membrane structural features such as packing defects and electrostatic interactions to αSyn binding across membrane phases remain unresolved.

Here, we investigated the binding behavior of N-terminally acetylated αSyn, the predominant physiological form of the protein^35–42^, as a function of membrane phase and charge using fluorescence microscopy and circular dichroism (CD) spectroscopy in combination with coarse-grained molecular dynamics (MD) simulations. Small unilamellar (SUVs) with defined lipid compositions were used to independently control membrane phase and charge. Under zwitterionic conditions and high curvature, αSyn preferentially binds to gel-phase membranes; however, this preference is reversed upon incorporation of anionic lipids, which selectively enhance binding in liquid-phase membranes. Coarse-grained MD simulations of flat and mechanically buckled lipid bilayers reveal that membrane phase governs curvature-dependent defect formation, providing a structural basis for these observations. These results further show that defect-mediated binding dominates in gel-phase membranes, whereas electrostatic interactions play a larger role in liquid-phase membranes. Finally, dynamic binding measurements demonstrate that membrane phase and charge regulate not only the extent of αSyn binding but also the stability and exchange behavior of the membrane-bound state. Together, these findings establish a framework in which membrane phase, curvature, and charge act cooperatively to regulate equilibrium partitioning, curvature sensitivity, and binding dynamics of αSyn.

## Results

### Impact of membrane phase and charge on αSyn binding to highly curved SUVs

To assess the impact of membrane phase and charge on αSyn binding, we utilized the lipid species shown in Figure 1A. Here, dioleoylphosphatidylcholine (DOPC) and dipalmitoylphosphatidylcholine (DPPC) represent lipids that form neutral liquid– and gel-phase membranes at room temperature, respectively, while dioleoylphosphatidylserine (DOPS) and dipalmitoylphosphatidylserine (DPPS) are their anionic counterparts. We used a previously developed fluorescence microscopy technique to visualize binding between αSyn and SUVs (Fig. 1B)^43; 44^. αSyn colocalization was observed for all four vesicle compositions, although the extent of colocalization varied with vesicle size, with the smallest vesicles consistently exhibiting detectable signal (Fig. 1C). When comparing binding between highly curved SUVs (50 nm diameter) primarily composed of either DOPC or DPPC, αSyn exhibited stronger binding to gel-phase DPPC membranes than liquid-phase DOPC membranes (Fig. 1D, Supplementary Figs. S1-S2). However, when anionic charge was introduced through inclusion of 25 mol% phosphatidylserine (PS) in membranes, there was a marked inversion for phase preference, with αSyn binding most strongly to DOPC/DOPS membranes (Fig. 1D, Supplementary Figs. S3-S4). When 25 mol% anionic phosphatidic acid (PA) was used instead of PS, binding was further enhanced in liquid-phase dioleoyl (DO) membranes but not in gel-phase dipalmitoyl (DP) membranes (Supplementary Fig. S5). The binding trends across PC and PC/PS membranes were mirrored with circular dichroism (CD) spectroscopy measurements using SUVs with approximately 70 nm diameter (Fig. 1E-F, Supplementary Fig. S6). Here, the characteristic α-helical spectra confirmed membrane insertion of αSyn’s NTD into membranes (Fig. 1E). These results reveal that electrostatic enhancement of αSyn binding is phase-dependent, with charge selectively amplifying association in liquid-phase membranes.

**Figure 1.**
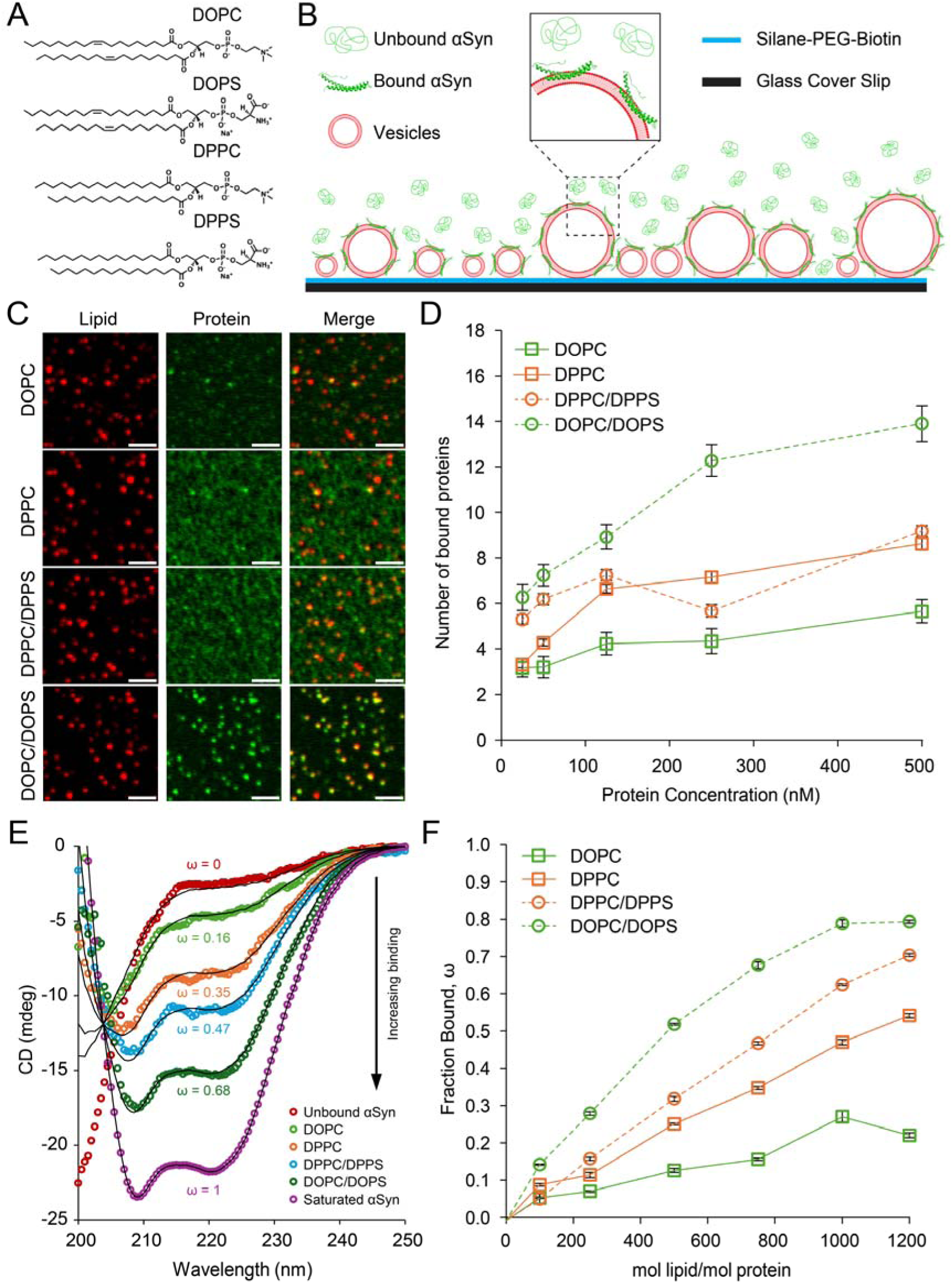
Headgroup charge inverts αSyn phase preference, favoring gel membranes when neutral and liquid membranes when anionic. (A) Chemical structures of liquid-phase (DOPC and DOPS) and gel-phase (DPPC and DPPS) lipids. (B) Schematic of the tethered vesicle assay. The schematic is not drawn to scale and is not intended to quantitatively reflect αSyn size or penetration depth. (C) Representative micrographs of tethered SUVs in the presence of 500 nM αSyn. Scale bars represent 2 μm. (D) Binding curves, determined through fluorescence microscopy, for αSyn association with SUVs with an average diameter of 50 nm. N = 604-3115 vesicles for each point on the binding curve. Error bars represent the 99% confidence interval of the mean. (E) Representative CD spectra demonstrating the extremes used for fitting ω, the fraction of αSyn in the bound state. The CD spectra correspond to 750:1 molar ratio of lipid:protein. Equation 1 was used for fitting and determining intermediate ω values. (F) Binding curves for unlabeled αSyn determined via CD spectroscopy. Error bars represent the standard error of the mean (SEM) for the fitting parameter, ω.

### Gel-phase membranes form faceted vesicles

Given αSyn’s enhanced binding to zwitterionic gel-phase membranes (Fig. 1D-F), we next examined whether membrane morphology differed between phases using cryo-electron microscopy (cryoEM) to visualize the morphology of SUVs that were extruded through 50 nm pores (Fig. 2A-B). Vesicle morphology was largely unaffected by headgroup charge. Liquid-phase DO vesicles displayed smooth, uniformly rounded morphologies (Fig. 2C) whereas gel-phase DP vesicles exhibited pronounced deviations from circularity, forming polygonal structures with regions of elevated local curvature (Fig. 2D). To quantify vesicle shape, circularity was calculated for each composition using the standard circularity metric (Equation 2), where a value of 1 corresponds to a perfect circle and lower values reflect deviations from a circle. (Fig. 2E). Liquid-phase vesicles composed of either DOPC or DOPC/DOPS exhibited circularity values closest to a perfect circle, whereas gel-phase vesicles showed reduced circularity scores (Fig. 2F). Collectively, these data establish that SUVs consisting of gel-phase membranes have discrete high-curvature regions that are not present in the liquid phase.

**Figure 2.**
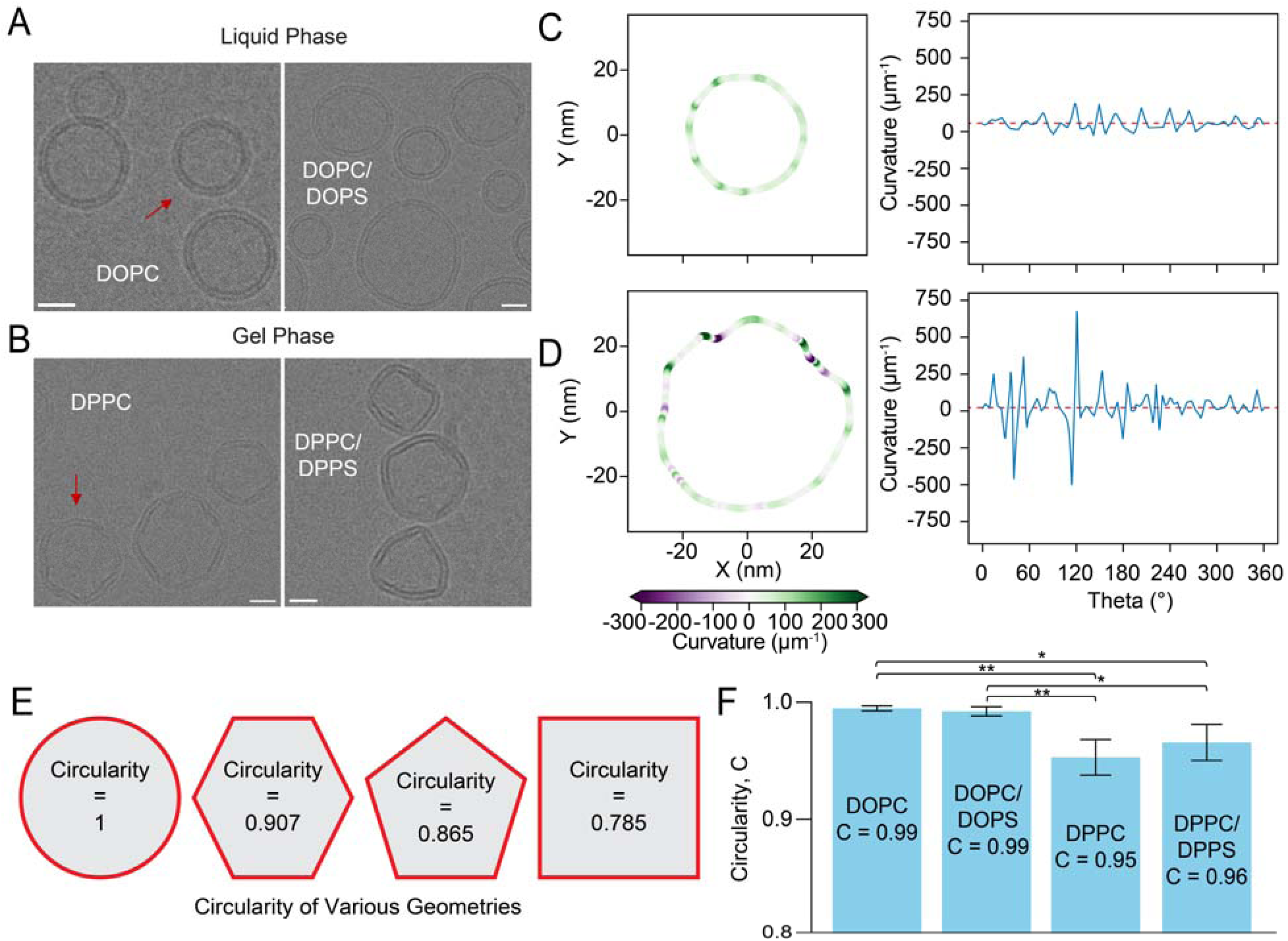
Gel-phase membranes adopt faceted vesicle morphologies, in contrast to spherical liquid-phase SUVs. CryoEM images of SUVs composed of (A) liquid-phase or (B) gel-phase lipid compositions. Red arrows indicate the vesicles analyzed in panels C and D. Scale bars indicate 20 nm. PC/PS mixtures were combined at a molar ratio of 3:1 PC:PS. (C) (left) Trace of the vesicle indicated by the red arrow in A, and (right) the corresponding curvature profile along the vesicle perimeter. (D) (left) Trace of the vesicle indicated by the red arrow in B, and (right) the corresponding curvature profile along the vesicle perimeter. Red-dashed line in the right panel of C and D corresponds to the average curvature of the analyzed vesicles. (E) Representative circularity values for various geometries. (F) Circularity values for each composition. Error bars represent the standard deviation (N = 3 measured vesicles). * corresponds to *P < 0.05* and ** corresponds to *P < 0.01* as determined by unpaired Student’s t-test.

### Curvature disproportionately enriches packing defects in gel-phase membranes

The enhanced association of αSyn with zwitterionic DPPC vesicles over DOPC vesicles (Fig. 1), together with the faceted morphology of gel-phase membranes (Fig. 2), suggests that localized high-curvature regions may create physical features that promote binding. We therefore asked whether bending gel-phase bilayers generates packing defects capable of explaining the observed phase-dependent binding preference. To quantify defect formation as a function of curvature and phase, we performed coarse-grained molecular dynamics simulations of flat (control) and mechanically buckled DOPC (liquid-phase) and DPPC (gel-phase) bilayers (Fig. 3). To facilitate the direct comparison between systems of varying lipid compositions, we matched the strain of each buckled system. The strains for buckled membrane systems were selected such that the membrane adopted curvatures that were comparable to those observed in cryoEM (Fig 2C-D, Supplementary Fig. S7). Representative side and top views of flat and buckled membranes are shown in Figures 3A-B. Defects, indicated by exposed acyl tail beads, in each of these membrane systems are shown in Figure 3C.

**Figure 3.**
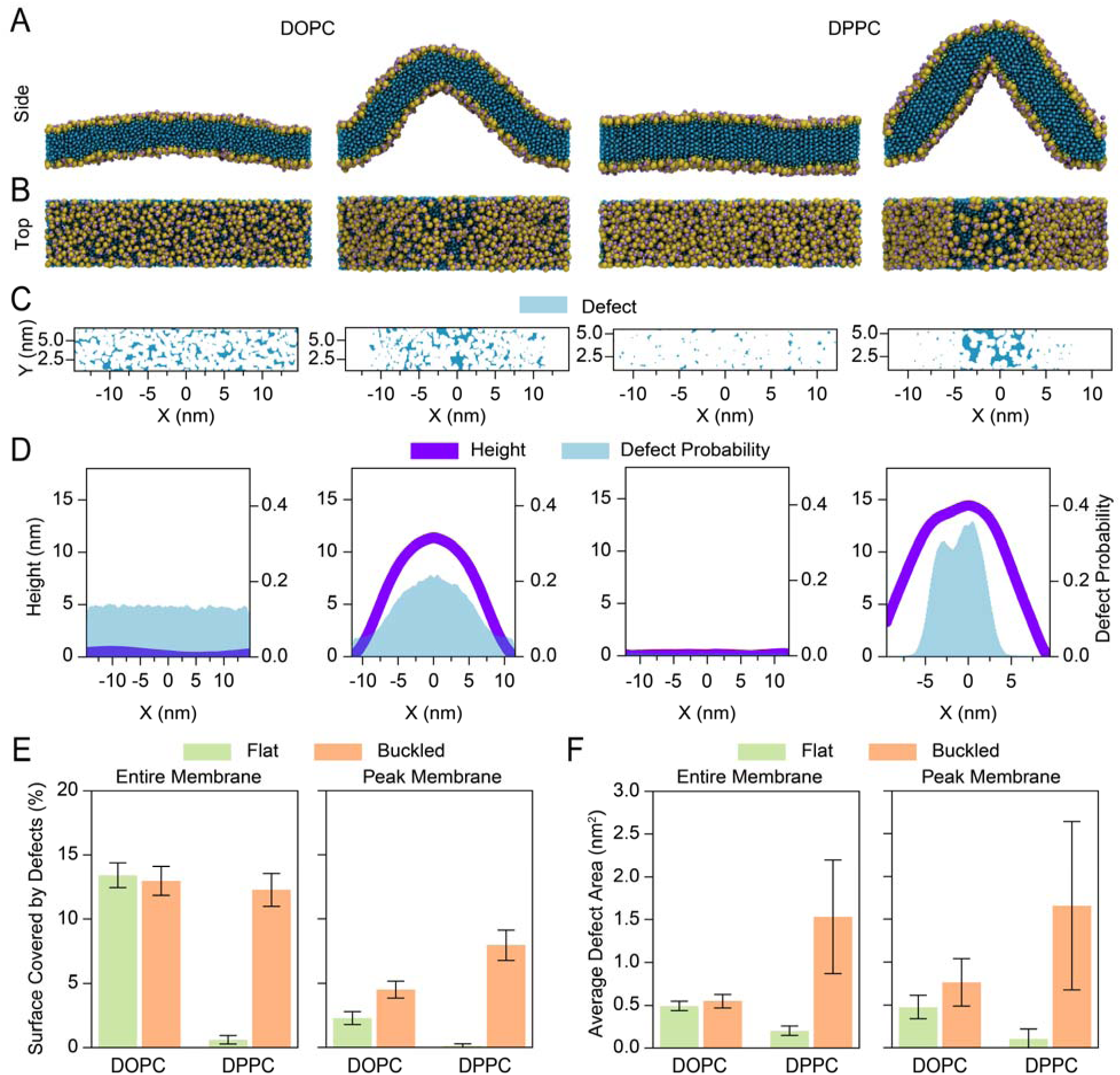
Buckling induces localized defect enrichment in DPPC but not DOPC membranes. Representative (A) side and (B) top views of flat and mechanically buckled coarse-grained DOPC (liquid-phase) and DPPC (gel-phase) bilayers. Yellow beads represent polar phosphate groups, green beads denote glycerol backbones, purple beads denote additional head group beads unique to each lipid, and blue beads indicate acyl chain groups identified as packing defects when exposed. (C) Lateral maps of defect locations for the representative bilayer above. (D) Spatial probability distribution of defects across the membrane. (E) Defect coverage and (F) average defect area across the entire membrane surface or restricted to the high-curvature crest region for flat and buckled configurations. Error bars in (E) and (F) represent the standard deviation calculated over all frames of the simulation.

To gain insight into how defect propensity changed with respect to curvature, we quantified the probability of finding a defect along the buckled direction (x-dimension) of the membrane (Fig. 3D). In liquid-phase DOPC membranes, defects were present in both flat and buckled configurations and remained uniformly distributed across the membrane surface. Imposing curvature produced little change in either total defect coverage or spatial organization (Fig 3D-F). In contrast, flat gel-phase DPPC membranes exhibited minimal defect content; however, upon buckling, defects increased markedly and became concentrated in the regions of highest curvature (Fig 3D-F).

Quantification of total defect coverage and defect size confirmed that in flat DOPC membranes, defects were larger and more prevalent compared to flat DPPC membranes (Fig. 3E-F, Supplementary Figs. S8-S9). Importantly, buckling did not significantly alter defect coverage in DOPC membranes, whereas DPPC membranes exhibited a substantial increase in defect size and defect coverage. When the analysis was restricted to the regions of highest curvature, this disparity became more pronounced: defect enrichment remained minimal in DOPC but increased sharply in DPPC (Fig. 3E-F). Incorporation of either PS or PA in the membranes resulted in the same qualitative trends where defect coverage and size dramatically increased when gel-phase membranes were buckled, but not when liquid-phase membranes were buckled (Supplementary Figs. S10-S11).

These simulations reveal strong curvature-defect coupling in gel-phase membranes that is largely absent in liquid-phase membranes. Although flat gel bilayers are defect-poor, membrane bending generates localized regions of elevated defect density. These findings provide a physical basis for how faceted, highly curved gel-phase vesicles can present defect-rich domains that promote αSyn association, consistent with the enhanced binding behavior observed for zwitterionic DPPC vesicles in Figure 1.

### Curvature sensitivity of αSyn is enhanced in gel-phase membranes and reduced by anionic charge

Since αSyn membrane curvature sensing is linked to membrane packing defects, the phase-dependent defect enrichment observed in Figure 3 suggests that curvature sensitivity should vary across membrane phases. To test this, we quantified protein partitioning as a function of vesicle diameter spanning approximately 30-150 nm (Fig. 4) using the tethered vesicle assay developed in Figure 1B. For liquid-phase DOPC and DOPC/DOPS membranes, αSyn binding increased monotonically with vesicle diameter, consistent with increasing available membrane surface area (Fig. 4A). In contrast, gel-phase DPPC and DPPC/DPPS membranes exhibited a non-monotonic binding profile, with maximal αSyn association observed near ∼50 nm vesicle diameter (Fig. 4B). Rather than scaling with surface area, binding peaked at intermediate curvature, indicating preferential association with smaller, more highly curved vesicles.

**Figure 4.**
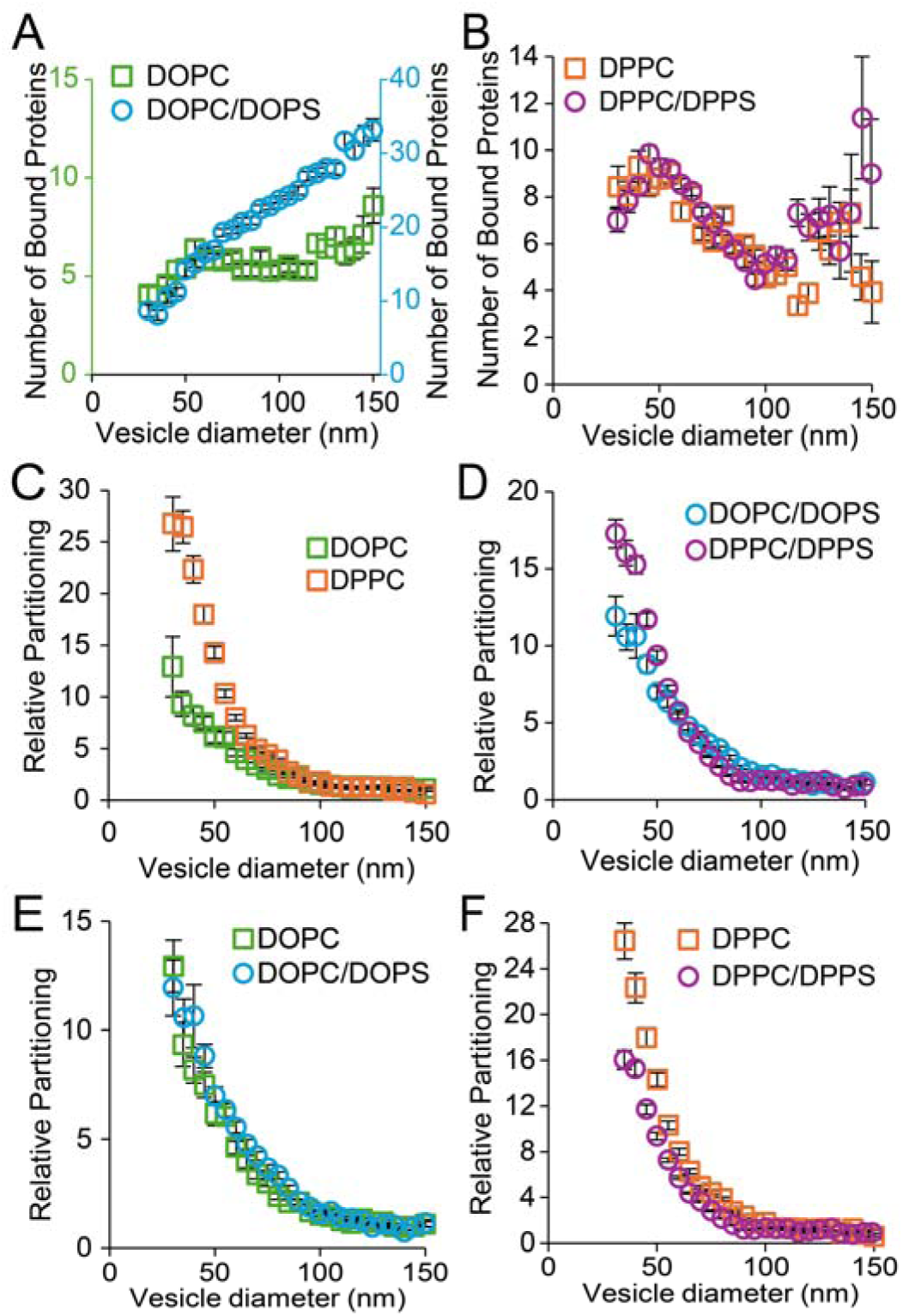
αSyn exhibits enhanced curvature sensitivity on gel-phase membranes. Membrane binding as a function of vesicle diameter for (A) liquid-phase and (B) gel-phase lipid compositions obtained via fluorescence microscopy. (C-F) Relative partitioning of αSyn to vesicles as a function of vesicle diameter. Binding density was normalized to the average protein density on vesicles with diameters between 130-150 nm. Error bars represent the 99% confidence interval of the mean for all panels. PC/PS mixtures were combined at a molar ratio of 3:1 PC:PS.

As the total protein bound increases with vesicle surface area, we directly assessed curvature sensitivity by normalizing protein density within each diameter bin to vesicles between 130-150 nm, which was reported as the relative partitioning of αSyn (Figs. 4C-F). αSyn displayed strong positive curvature sensitivity, with partitioning increasing as curvature increased. Under zwitterionic conditions, αSyn displayed substantially greater curvature sensitivity on DPPC vesicles than on DOPC vesicles, confirming enhanced curvature sensitivity in gel-phase membranes (Fig. 4C). When anionic PS headgroups were present, curvature sensitivity remained greater in gel-phase membranes relative to liquid-phase membranes, but the magnitude of the difference was reduced (Fig. 4D).

To evaluate the effect of charge within each phase independently, we compared the curvature sensitivity of αSyn bound to zwitterionic and anionic membranes of identical phases. In liquid-phase DO membranes, incorporation of anionic PS headgroups produced minimal change in curvature sensitivity (Fig. 4E), while in gel-phase DP membranes, addition of PS significantly reduced curvature-dependent enrichment (Fig. 4F). Interestingly, the use of PA headgroups instead of PS headgroups substantially reduced curvature preference in both phases (Supplementary Fig. S12). These results indicate that αSyn curvature sensitivity is stronger in gel-phase membranes and that anionic charge more strongly modulates curvature sensitivity in this phase, with the exception of PA lipids, which diminished curvature sensitivity in both gel and liquid phases.

### Membrane charge modulates αSyn binding dynamics across membrane phases

Because αSyn’s function at the synapse depends on dynamic membrane association with synaptic vesicles^7^, we next asked how membrane phase and charge regulate the dynamic binding behavior of αSyn. To probe binding dynamics, we performed fluorescence recovery after photobleaching (FRAP). In these FRAP experiments, a defined region of interest containing fluorescently labeled, membrane-bound αSyn was photobleached, and fluorescence recovery was monitored over time (Fig. 5A). Recovery curves were fit using Equation 5 to extract the mobile fraction (*x_m_*) and the dissociation rate constant of the mobile fraction (*k_off_*) (Fig. 5B). The fitted off-rates for the immobile fractions (k_off,i_) were 0 in all conditions, consistent with an immobile population that does not exchange on the timescale of our FRAP measurements; therefore, k_off,i_ was not plotted.

**Figure 5.**
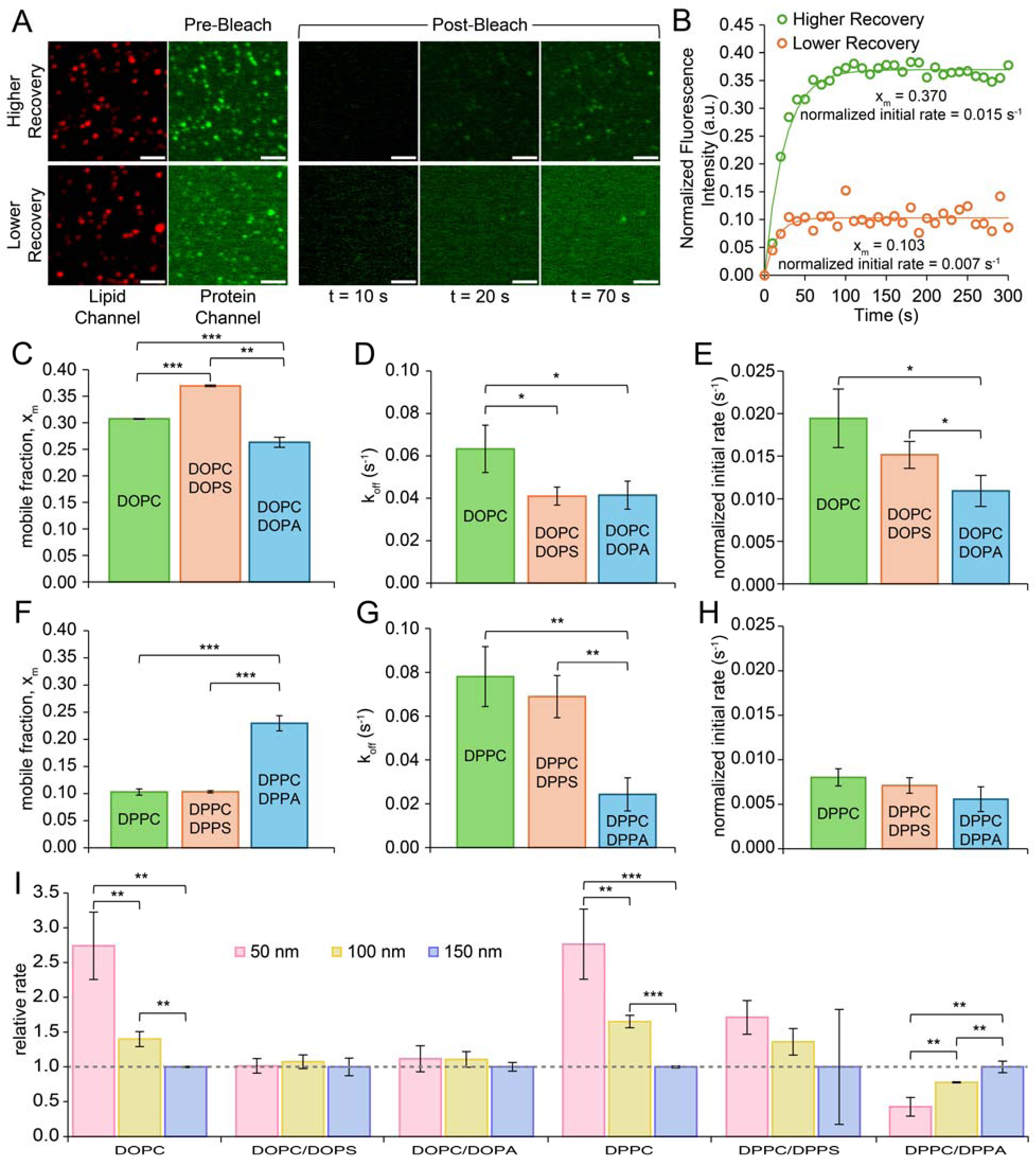
Anionic lipids reduce αSyn mobility and slow FRAP kinetics on liquid– and gel-phase membranes. (A) Representative fluorescence images of αSyn bound to lipid vesicles before and after photobleaching. SUVs were incubated with 1 μM protein. “Higher Recovery” corresponds to 3:1 DOPC:DOPS vesicles, while “Lower Recovery” corresponds to 3:1 DPPC:DPPS vesicles. Scale bars represent 2 μm. (B) Representative fluorescence recovery curves with fits determined by using Equation 5. (C, F) Fraction of αSyn that is mobile on the membrane surface after photobleaching (x_m_). (D, G) Dissociation rate constant (k_off_) of αSyn from the membrane after photobleaching. (E, H) Normalized initial rate of αSyn dissociation, defined as the product of x_m_ and k_off_. (I) Initial rate of αSyn dissociation relative to the rate on vesicles with 150 nm diameter. Error bars represent the standard deviation (n = 3 FRAP fields of view, each containing 35-768 vesicles). Dashed line serves as a visual guide for the condition to which all measurements were normalized. * corresponds to *P < 0.05*, ** corresponds to *P < 0.01*, and *** corresponds to *P < 0.001* as determined by unpaired Student’s t-test. All measurements were performed at room temperature, approximately 20°C.

All FRAP analyses in Figures 5B-H were performed on vesicle populations with an average diameter of 50 nm. Although x_m_ and k_off_ varied modestly across conditions (Figs. 5C,D,F,G), the clearest trend emerged from the normalized initial rate, x_m_k_off_, which represents the overall binding recovery kinetics (Figs. 5E,H). These overall recovery rates were higher on liquid-phase DO membranes than on gel-phase DP membranes, indicating faster exchange on liquid-phase membranes and slower exchange on gel-phase membranes. Within each phase, incorporation of anionic lipids reduced the normalized initial rate, with the strongest reduction observed for PA-containing membranes. This reduction was statistically significant in liquid-phase membranes but not in gel-phase membranes, indicating that anionic lipids more strongly suppress αSyn exchange dynamics in fluid membranes.

To determine whether αSyn exchange dynamics were impacted by membrane curvature, we compared normalized initial rates for vesicles with average diameters of 50, 100, and 150 nm (Fig. 5I). Here, the relative rate is the normalized initial rate for a specific condition divided by the corresponding rate on 150 nm vesicles. On zwitterionic DOPC and DPPC membranes, αSyn exhibited positive curvature sensitivity, with greater recovery rates on smaller, more highly curved vesicles (Fig. 5I, Supplementary Figs. S13-S14). Introduction of anionic charge to liquid-phase DO membranes nearly abolished this dynamic, positive curvature sensitivity. In the gel phase, positive curvature sensitivity was diminished with the addition of PS and completely inverted with the addition of PA, eliciting a negative curvature sensitivity response. Together, these results demonstrate that anionic lipids reshape curvature-dependent αSyn exchange kinetics in a phase-dependent manner, attenuating curvature sensitivity in liquid phase membranes while diminishing or reversing it in gel-phase membranes.

### αSyn binding across gel-liquid phase transitions

So far, we have examined αSyn binding within individual lipid phases. However, lipid membranes are often not restricted to a single phase; they can transition between phases or exhibit coexistence of multiple phases. Biological membranes are highly dynamic and heterogeneous^45^ and αSyn binding has been shown to change sharply near phase boundaries^24^. Therefore, we examined how membrane association evolves across gel-liquid phase transitions and how anionic charge modulates binding in this regime. To quantify αSyn binding across membrane phase transitions, we performed CD temperature ramp experiments on gel-phase DP membranes. Representative CD spectra for αSyn bound to DPPC vesicles are shown in Figure 6A, demonstrating progressive strengthening of the α-helical signature as temperature increases toward the melting temperature (T_m_). The full temperature-dependent evolution of the CD spectra is shown in Figure 6B, which reveals a pronounced increase in helical content approaching the T_m_, followed by a rapid loss of signal above the phase transition. CD spectra were fit using Equation 1 to extract the fraction of membrane-bound αSyn at each temperature. Fluorescence anisotropy measurements were used to determine the melting temperatures of each composition and the width of the phase transition window (Supplementary Figs. S15-S16, See methods). For pure DPPC membranes, αSyn binding increased sharply upon heating from ∼30 to just below T_m_ (∼40°C), reached a maximum near the phase boundary, and then decreased rapidly once the temperature transitioned fully into the liquid phase (Fig. 6C). Above the T_m_, αSyn remained largely unbound. A similar pre-transition increase in binding was observed for DPPC/DPPS membranes (Fig. 6D). However, in contrast to pure DPPC, significant αSyn binding persisted across and beyond the melting transition. Although binding reached a maximum near the T_m_ (∼41°C), measurable membrane association was maintained until 50°C before complete dissociation occurred. Thus, incorporation of PS broadened the temperature window over which αSyn remained membrane-bound, stabilizing association above the gel-liquid phase boundary.

**Figure 6.**
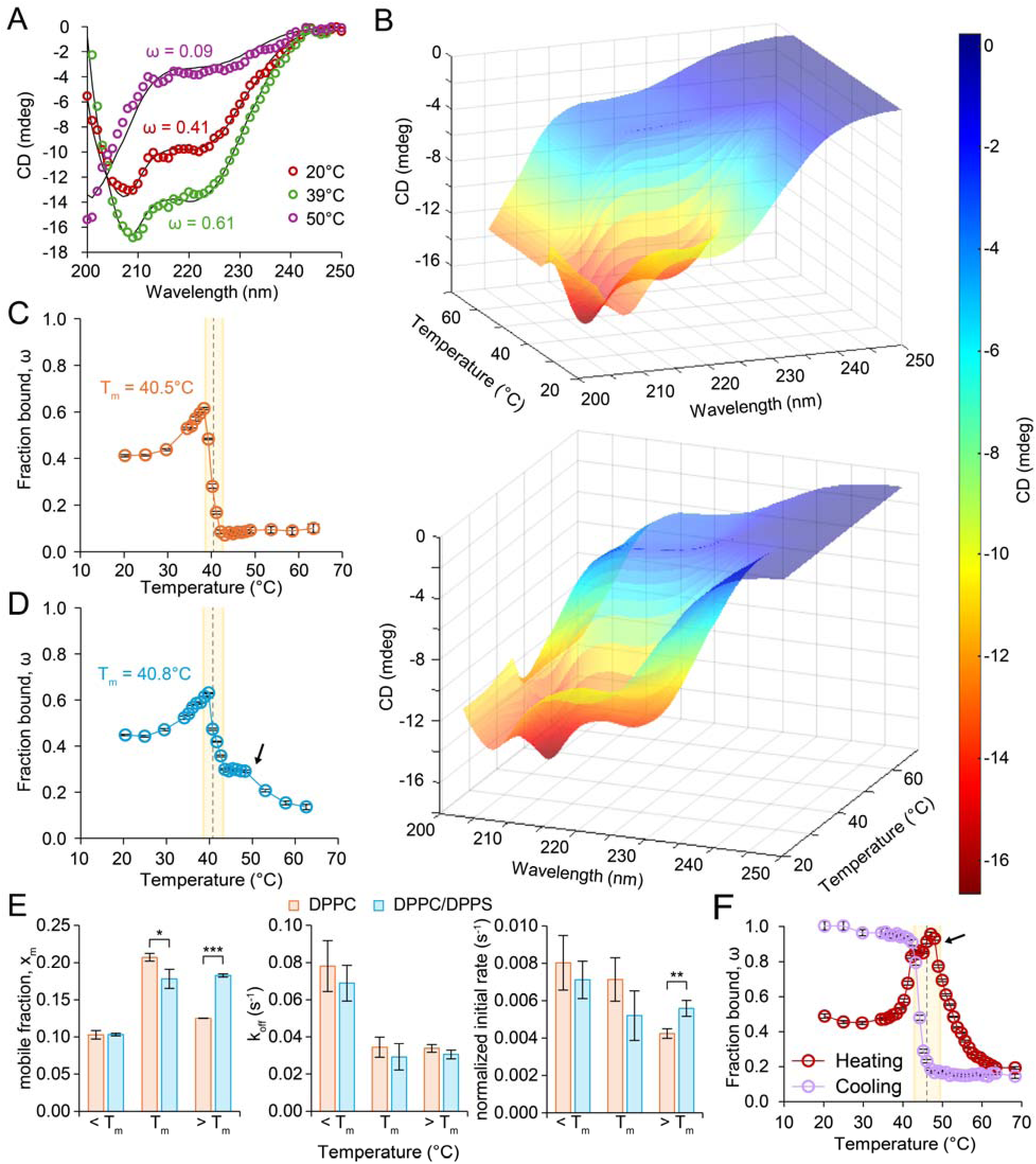
Anionic lipids stabilize αSyn binding across the gel-liquid phase transition. (A) Representative CD spectra demonstrating temperature-dependent changes in αSyn secondary structure upon binding to DPPC vesicles, with reduced binding above the lipid melting temperature T_m_. (B) Representative three-dimensional CD spectra illustrating changes in αSyn binding as temperature is increased from 20°C to 63°C in the presence of DPPC SUVs. (C) Temperature-dependent binding of αSyn to DPPC SUVs quantified from CD spectroscopy. Error bars represent the SEM for the regression parameter ω at each temperature. (D) Temperature-dependent binding of αSyn to 3:1 DPPC:DPPS SUVs quantified from CD spectroscopy. Error bars represent the SEM for the regression parameter ω at each temperature. (E) (left) Mobile fraction (X*_m_*), (middle) dissociation rate constant (k*_off_*), and (right) normalized initial rate (*x_m_*,k*_off_*) of αSyn dissociation on 100% DPPC and 3:1 DPPC:DPPS vesicles below, at, and above T_m_. Error bar represents standard deviation (n = 3). * corresponds to *P < 0.05*, ** corresponds to *P < 0.01*, and *** corresponds to *P < 0.001* as determined by unpaired Student’s t-test. (F) Temperature-dependent binding of αSyn to 3:1 DPPC/DPPA SUVs quantified from CD spectroscopy. Error bars represent the SEM for the regression parameter ω at each temperature. Shaded regions in C, D, and F indicate the phase boundary, and dashed lines denote T_m_. Black arrows in D and F mark the temperature at which αSyn binding destabilizes.

To determine whether exchange dynamics were similarly altered near the phase transition, we performed FRAP measurements below, at, and above T_m_ for DPPC and DPPC/DPPS membranes (Fig. 6E). For both compositions, x_m_ increased at T_m_, indicating that a larger fraction of membrane-bound αSyn becomes exchangeable at the phase boundary. Above T_m_, however, x_m_ decreased for DPPC but remained elevated for DPPC/DPPS, demonstrating that PS sustains a larger exchangeable population in the liquid phase. In contrast, k_off_ decreased at T_m_ and remained low above T_m_ for both compositions, indicating slower dissociation kinetics across the transition. Thus, the presence of PS does not accelerate molecular turnover; rather, it preserves a greater fraction of αSyn in a dynamically exchanging state while maintaining low dissociation kinetics. Consistent with these combined effects, the normalized initial recovery rate (x_m_k_off_) diverged most clearly above T_m_, where DPPC/DPPS exhibited higher recovery dynamics than DPPC. Together, these data show that PS stabilizes αSyn binding across the phase transition by retaining slow off-rates while preventing the collapse of the exchangeable population that occurs in pure DPPC membranes.

We next performed the CD temperature ramp measurements on DPPC/DPPA membranes (Fig. 6F). Upon heating, αSyn binding saturated immediately prior to the melting transition (∼47°C) and remained elevated until approximately 50°C, where binding eventually decreased. During cooling, αSyn rebounded sharply upon crossing T_m_, with saturated binding between 20-40. Notably, neither DPPC nor DPPC/DPPS membranes exhibited comparably strong hysteresis (Supplementary Figs. S17, S19). DOPC and DOPC/DOPS controls, which do not undergo phase transitions over this temperature range, similarly lacked strong hysteresis (Supplementary Figs. S18, S20). Together, these observations indicate that PA confers even greater stabilization of αSyn binding across the gel-liquid phase transition than PS. Due to the elevated melting temperature of DPPC/DPPA membranes and reduced αSyn stability at the required temperatures, FRAP measurements could not be reliably performed for this composition above its T_m_, precluding quantitative analysis of exchange dynamics in this regime.

## Discussion

In this study, our results highlight the phase-dependent behavior of αSyn-membrane interactions. On highly curved vesicles composed of zwitterionic lipids, αSyn binds more strongly to gel-phase than liquid-phase membranes, indicating that affinity in this curvature regime is governed by lipid tail order. Consistent with this result, αSyn exhibits greater curvature sensitivity on gel-phase membranes, demonstrating that membrane phase is a factor influencing the relationship between membrane geometry (i.e., curvature) and αSyn binding. CryoEM and simulations reveal that highly curved gel-phase vesicles adopt faceted morphologies, consistent with previous reports^46–48^. In this faceted geometry, relatively flat faces are separated by sharply curved ridges or vertices. Lipids in the gel phase are well-packed, forming flat membrane configurations. When the gel-phase membrane is forced to conform to high curvature (e.g., that imposed by an SUV), the membrane will ‘crack’, concentrating the lipid packing defects to these points. Along with the disruption in acyl tail nematic order, on the surface of the membrane, at the vertices of faceted gel-phase SUVs, surface packing defects are also concentrated. With sufficient faceting, gel-phase vesicles can present greater overall surface packing defect content than liquid-phase vesicles, explaining the enhanced binding observed in this high curvature regime. This framework also accounts for the non-monotonic binding profile in gel-phase vesicles, with maximal αSyn binding near ∼50 nm diameter (Fig. 4B). This binding preference may reflect a reduction in the fractional area of defect-rich facet boundaries as vesicle diameter increases. It is further plausible that defect abundance can explain the rise in protein binding near the phase transition temperature of DPPC-containing membranes during heating. The co-existence of phases leads to increased phase boundaries which foster surface packing defects.

Incorporation of anionic lipids dramatically enhanced binding in liquid-phase membranes but had only modest impact on binding to gel-phase membranes, indicating that the contributions of electrostatics to binding are attenuated by acyl chain ordering. Electrostatic enhancement of αSyn binding is well established, arising from interactions between the positively charged N-terminal region and anionic headgroup^19–23; 26; 44; 49–51^. We expect these interactions to depend on lipid mobility. αSyn binding likely involves local lipid rearrangements, such as lateral redistribution of anionic lipids and accommodation of the amphipathic helix within the bilayer, which are processes that are readily facilitated in fluid membranes^52–55^. In gel-phase membranes, reduced lipid mobility limits such reorganization, constraining the extent to which electrostatics can stabilize αSyn insertion. Thus, while defect enrichment explains enhanced binding to zwitterionic DPPC relative to DOPC, we suggest that electrostatic interactions dominate binding enhancement in liquid-phase membranes. The stronger enhancement of binding observed with DOPA compared to DOPS further suggests that anionic charge alone does not fully define this effect; headgroup size and steric accessibility likely influence how effectively αSyn engages the membrane interface.

This distinction also explains why anionic lipids reduce curvature sensitivity in gel-phase membranes. In gel-phase vesicles, binding to highly curved membranes is largely governed by curvature-induced defect enrichment. When this structural contribution is large, additional electrostatic stabilization provides only a modest incremental effect, particularly given the restricted lipid mobility. In contrast, larger gel-phase vesicles contain fewer defects and exhibit weaker baseline binding (Fig. 4B), making electrostatic contributions more apparent and compressing curvature-dependent differences (Fig. 4F). This interpretation is consistent with the two-stage binding model proposed by Bartels *et al.*^25^, in which initial N-terminal membrane recognition can be supported by electrostatics or packing defects, whereas subsequent cooperative helix formation likely depends on the membrane’s ability to accommodate local lipid rearrangement. In liquid-phase membranes, where curvature has a smaller impact on defect structure and lipid rearrangement is readily accommodated, electrostatic interactions enhance binding more uniformly across vesicle sizes without substantially altering curvature preference. Notably, our coarse-grain simulations did not capture an increase in defect abundance upon incorporation of anionic lipids in liquid-phase membranes, despite the clear enhancement in αSyn binding. Prior atomistic simulations revealed modest but detectable increases in defect abundance for DOPC/DOPS bilayers relative to DOPC^44^. This may be due to differences between the atomistic and coarse-grained representations and, in particular, the choices of Van der Waals radii for each representation. A similar limitation likely applies to DOPC/DOPA membranes, where the smaller PA headgroup would be expected to reduce steric shielding at the interface and potentially promote local defect formation, but this effect was not resolved here. Importantly, the simulations robustly capture the large defect contrast between liquid– and gel-phase membranes, indicating that the dominant phase-dependent trends are well resolved. These findings highlight that defect abundance alone cannot fully account for the observed charge-dependent binding trends. Consistent with the established role of electrostatics in αSyn membrane binding^19;56;57^, electrostatic contributions, which are not explicitly captured in our modeling framework, likely underlie the enhanced association observed in liquid-phase membranes.

Consistent with this interpretation, dynamic exchange measurements further reveal how membrane phase and charge influence not only the extent of αSyn binding, but also the stability of the bound state. αSyn exhibits slower exchange kinetics and reduced mobile fractions on gel-phase membranes, consistent with a rigid environment that stabilizes bound states. In the liquid-phase membranes, αSyn exchange rates decreased directionally with incorporation of PS and PA lipids, suggesting electrostatic stabilization of binding. In the absence of charge, exchange dynamics exhibit positive curvature dependence in both phases, with faster exchange on highly curved vesicles, consistent with increased defect availability and acyl chain splaying. Incorporation of anionic lipids dampens this curvature dependence, with the most pronounced effect observed for DPPC/DPPA, where dynamics were inverted to reflect negative curvature sensitivity. The stronger effect of PA relative to PS suggests that headgroup structure, not charge alone, contributes to this behavior. In highly curved gel-phase membranes, the smaller PA headgroup may permit tighter αSyn engagement with defect-rich regions, converting highly curved vesicles from rapidly exchanging surfaces into more kinetically stabilized binding environments. In contrast, the larger PS headgroup may provide electrostatic stabilization without enabling the same degree of interfacial engagement, thereby dampening curvature-dependent exchange without fully inverting it. Together, these results show that membrane phase and charge influence not only the extent of binding but also the stability of the bound state, with gel-phase membranes dominated by structural defect effects and liquid-phase membranes by electrostatic stabilization.

Finally, anionic lipids stabilize αSyn binding across gel-liquid phase transitions. Binding increases sharply near T_m_, potentially reflecting the formation of a ripple phase^58–62^ or gel-phase liquid coexistence^63; 64^, both of which introduce additional defect-rich environments. In PS-containing membranes, electrostatic recruitment may become especially effective near and above T_m_, where transition-associated defects provide insertion sites and increased lipid mobility lowers the energetic cost of accommodating the amphipathic helix. Consistent with this interpretation, membrane exchange dynamics remain stabilized above T_m_ for DPPC/DPPS relative to DPPC. This effect is further enhanced in DPPC/DPPA, where the smaller PA headgroup may promote stronger interfacial engagement, producing pronounced hysteresis with stronger binding during cooling than heating. This behavior may arise from differences in membrane state during initial αSyn engagement, where liquid-phase membranes during cooling permit deeper insertion and stronger electrostatic coupling prior to defect formation. The reduced binding near ∼50 suggests the onset of thermal destabilization of the αSyn-membrane complex. Taken together, our findings demonstrate that membrane phase, curvature, and charge jointly regulate αSyn binding through distinct yet interconnected mechanisms. In gel-phase membranes, curvature-induced faceting and defect enrichment dominate both equilibrium and dynamic behavior, producing strong curvature sensitivity that can be partially modulated by electrostatics. In liquid-phase membranes, lipid mobility enables electrostatic interactions to strengthen and prolong binding without substantially altering curvature preference. Thus, structural defects govern phase– and curvature-dependent behavior, while electrostatics tune binding strength and stability in a mobility-dependent manner.

## Conclusion

In conclusion, this study demonstrates that αSyn binding to lipid membranes is governed by the combined effects of membrane phase and charge rather than either property in isolation. Under equilibrium conditions, αSyn preferentially associates with zwitterionic gel-phase membranes; however, the introduction of anionic lipids inverts this trend, promoting enhanced binding to liquid-phase membranes. Under dynamic conditions, αSyn exhibits reduced mobility on gel-phase membranes relative to liquid-phase membranes, with membrane charge further stabilizing membrane association and diminishing curvature-dependent exchange behavior. Across gel-liquid phase transitions, anionic lipids extend the temperature range over which αSyn remains membrane-bound and modulates its exchange dynamics.

These findings emphasize that αSyn-membrane interactions cannot be understood through membrane phase or charge alone but instead arise from their interplay in a curvature-dependent context. In particular, curvature-dependent defect formation dominates αSyn binding behavior in gel-phase membranes, while electrostatic interactions play a larger role in stabilizing association in liquid-phase membranes. Rather than reflecting a universal preference for a particular membrane phase, our results indicate that αSyn binding depends sensitively on membrane curvature, phase, and composition.

More broadly, these results provide a framework for understanding how changes in membrane physical properties influence αSyn association. In biological membranes, lipid composition, curvature, and phase behavior are dynamically regulated and can shift with aging and disease. Our findings suggest that such changes may alter αSyn binding by modifying defect landscapes and electrostatic interactions, thereby influencing the stability and persistence of membrane-associated states. Future studies incorporating these additional complexities may help clarify how alterations in membrane properties contribute to αSyn-associated pathologies.

## Materials and Methods

### Materials

1,2-dioleoyl-sn-glycero-3-phosphocholine (DOPC), 1,2-dioleoyl-sn-glycero-3-phospho-L-serine (sodium salt) (DOPS), 1,2-dipalmitoyl-sn-glycero-3-phosphocholine (DPPC), 1,2-dipalmitoyl-sn-glycero-3-phospho-L-serine (sodium salt) (DPPS), 1,2-dipalmitoyl-sn-glycero-3-phophate (sodium salt) (DPPA), and 1,2-dioleoyl-sn-glycero-3-phosphate (sodium salt) (DOPA) were purchased from Avanti Polar Lipids. Isopropyl-beta-D-thiogalactoside (IPTG) was purchased from Gold Biotechnology. Tris(2-carboxyethyl) phosphine hydrochloride (TCEP), phenylmethanesulfonyl fluoride (PMSF), sodium phosphate monobasic, sodium phosphate dibasic, and β-mercaptoethanol (BME) were purchased from Sigma-Aldrich. 4-(2-hydroxyethyl)-1-piparazineethanesulphonic acid (HEPES) and tris(hydroxymethyl)aminomethane (Tris) were purchased from Fisher Scientific. ATTO 647N-labeled 1,2-Dipalmitoyl-sn-glycero-3-phosphoethanolamine (DPPE) and ATTO 488-maleimide were purchased from ATTO-TEC. Biotin-PEG-Silane (5000 MW) and mPEG-Silane (5000 MW) were purchased from Laysan Bio, Inc. Dipalmitoyl-decaethylene glycol-biotin (DP-EG15-biotin) was provided by D. Sasaki from Sandia National Laboratories, Livermore, CA^65^.

### Expression Purification of α-synuclein

A detailed protocol for expressing and purifying α-synuclein has been described previously^44^. Briefly, the plasmid encoding the Y136C α-synuclein variant, cloned into the pRK172 backbone (obtained from Jennifer C. Lee (National Institutes of Health, Bethesda, MD)), was introduced into BL21 (DE3) E. coli cells that had been previously transformed with the pNatB plasmid to enable N-terminal acetylation of α-synuclein^66^. Cultures were grown in 2xTY medium at 37 °C to an optical density at 600 nm of approximately 0.6, then induced by the addition of 1 mM IPTG for 4 hours. Following induction, cells were harvested by centrifugation and resuspended in lysis buffer containing 25 mM Tris, 20 mM NaCl, 1 mM PMSF, 10 mM β-mercaptoethanol, and a Roche protease inhibitor cocktail (Sigma-Aldrich) (pH 8.0). Cell disruption was carried out by sonication on ice at 65% amplitude for 15 minutes. DNase I (Thermo Fisher Scientific) (30 µL per 10 mL of lysate) was then added and incubated at 37 °C for 30 minutes. The lysate was acidified to pH 3.5, and insoluble material was removed by centrifugation. The clarified supernatant was subsequently neutralized to pH 7.0, followed by ammonium sulfate (Sigma Aldrich) precipitation to 50% saturation. After centrifugation, the resulting protein pellet was dissolved in 20 mM Tris (pH 8.0) and applied to a 5 mL HiTrap Q FF anion-exchange column (Cytiva). Protein elution was achieved using 20 mM Tris, 1 M NaCl, and 10 mM β-mercaptoethanol (pH 8.0). Eluted fractions were concentrated using a 3 kDa molecular weight cutoff centrifugal filter (Sigma-Aldrich) and further purified by size-exclusion chromatography on a Superdex 200 10/300 column (Cytiva) equilibrated in 25 mM HEPES, 150 mM NaCl, and 0.5 mM TCEP (pH 7.4). Fractions corresponding to monomeric α-synuclein were pooled and concentrated, and final protein concentrations were determined by UV–visible absorbance.

### Labeling α-synuclein

Fluorescence labeling of α-synuclein was performed as previously described^44^. First, a sixfold molar excess of ATTO 488-maleimide was combined with Y136C α-synuclein. The reaction was performed covered and overnight at 4 °C. Unreacted dye was then separated from labeled protein by size-exclusion chromatography using a Superdex 75 10/300 GL column (Cytiva). UV–visible spectroscopy confirmed labeling efficiency to be approximately 1:1 protein-to-dye.

### SUV Preparation

Lipid aliquots were thawed from –80 and combined in the following molar percentages for use in the tethered vesicles assays:

- 99% DOPC, 0.5% DP-EG15-biotin, 0.5% ATTO 647N
- 74% DOPC, 25% DOPS, 0.5% DP-EG15-biotin, 0.5% ATTO 647N
- 74% DOPC, 25% DOPA, 0.5% DP-EG15-biotin, 0.5% ATTO 647N
- 99% DPPC, 0.5% DP-EG15-biotin, 0.5% ATTO 647N
- 74% DPPC, 25% DPPS, 0.5% DP-EG15-biotin, 0.5% ATTO 647N
- 74% DPPC, 25% DPPA, 0.5% DP-EG15-biotin, 0.5% ATTO 647N

The following molar percentages were used for vesicle preparation in CD spectroscopy experiments:

- 100% DOPC
- 75% DOPC, 25% DOPS
- 100% DPPC
- 75% DPPC, 25% DPPS
- 75% DPPC, 25% DPPA

After combining the lipid mixtures, the chloroform was evaporated with a nitrogen stream. Lipid films were dried for a minimum of 2 hours before rehydration using 20 mM sodium phosphate and 150 mM NaCl (pH 7.0) buffer (Working Buffer). After 15 minutes of hydration, the lipid suspensions were subjected to 6 freeze/thaw cycles by submerging the solution in liquid nitrogen for 1 minute and thawing at 50-70 for 5 minutes. The SUVs were then extruded through a 50 nm polycarbonate membrane (Whatman) at room temperature for the DO lipids and heated above the melting temperature for the DP lipids. The average diameter of vesicles used for CD spec across all compositions was 70.4 ± 3.5 nm, determined by dynamic light scattering.

### Slide Passivation and Vesicle Tethering

Glass coverslips were cleaned and treated as previously described^44; 67; 68^. Glass-bottom dishes were cleaned by adding enough heated KOH/peroxide and HCl/peroxide solutions to cover the bottom of the dish. The surfaces were then passivated by treatment with a PEG-silane solution (10 mg/mL in isopropanol) consisting of 95% methoxy-PEG-silane and 5% biotin-PEG-silane. Glacial acetic acid was added to the silane mixture at a final concentration of 1% (v/v) to catalyze silane coupling to surface hydroxyl groups. 100 µL of the solution was applied to each cleaned dish, evenly distributed, and incubated at 50 °C for 1 hour. Before use, excess PEG-silane was removed by rinsing with deionized water and dried using compressed air. Prepared slides and dishes were stored dry at ambient conditions and used within one week. 0.8 mm thick silicone gaskets (Grace Bio-Labs) with 5 mm diameter holes were placed on the cleaned glasses to create imaging wells. To tether the vesicles, 30 μl of NeutrAvidin at 3.33 μM was added to each well and incubated for 20 minutes. Excess NeutraAvidin was removed by rinsing with the Working Buffer. Prepared vesicles with 1 μM bovine serum albumin (BSA) were then added to the wells at 1 μM and incubated for 10 minutes. Excess, unbound vesicles were rinsed with the Working Buffer. BSA was added to limit non-specific adsorption of lipid species to the treated glass surface. The proteins were then added to the wells at specific concentrations and imaged immediately.

### Confocal Microscopy

Fluorescence imaging was carried out using a laser scanning confocal microscope (Leica Stellaris 5). Labeled α-synuclein was excited using a 488 nm laser with emission signals collected over a window of 493-558 nm. Labeled SUVs were excited with a 638 nm laser and emission signals were collected over a window of 643–749 nm. A Leica HC PL APO 63× oil-immersion objective with a numerical aperture of 1.4 was used to acquire images. The optical zoom was adjusted to yield a pixel dimension of 70 nm × 70 nm, and all scans were recorded at a line frequency of 400 Hz.

For fluorescence recovery after photobleaching (FRAP) measurements, the image acquisition sequence consisted of a pre-bleaching imaging step, followed by photobleaching the region of interest (ROI), and then a post-bleaching step. For pre-bleaching and post-bleaching, the laser power was set to 4%. The bleaching of the ROI was performed at 100% laser power. Pre-bleaching was carried out by acquiring 3 images of the ROI (total time, 2 s), followed by bleaching with 20-50 bleach frames (total time, 13-33 s). Finally, 30 post-bleach images were acquired every 10 s. For heated FRAPs, samples were placed inside an H301-MINI stage incubator (Okolab), and the system was heated using Okolab Nano. To prevent evaporation of the samples, a humidifier was used during the experiments.

### Image Processing for Confocal Microscopy

Detailed image analysis has been described previously^44^. In brief, images of diffraction-limited fluorescent puncta corresponding to labeled lipids and proteins were identified and analyzed using the open-source software package cmeAnalysis^69^. Using the lipid-channel as the primary detection channel, our analysis only included puncta that were consistently detected across three consecutive imaging frames within a given field of view. Additionally, our analysis only included diffraction-limited vesicles with diameters less than 250 nm. Protein binding was quantified by assessing colocalized fluorescence signal in the protein-channel on all vesicles.

SUV diameters were determined by obtaining fluorescence intensity distributions representative of the vesicle population, then determining a scaling factor by aligning the square root of the vesicle fluorescence intensity distribution with the corresponding diameter distribution measured by dynamic light scattering (Supplementary Fig. S21A). For curvature-dependent analysis directly comparing 50 nm, 100 nm, and 150 nm diameter vesicles, bins were defined as follows: 40-60 nm for the 50 nm bin, 90-110 nm for the 100 nm bin, and 140-160 nm for the 150 nm bin.

The number of bound proteins was determined via calibration experiments where ATTO 488-labeled α-synuclein was absorbed on glass coverslips and fluorescence intensity distributions were obtained. The peak intensity of the distribution was taken as the signal corresponding to a single labeled protein (Supplementary Fig. S21B). For fluorescence imaging experiments with different detector gains, this single molecule intensity scaled accordingly.

### Circular Dichroism Spectroscopy

Single temperature circular dichroism scans were performed as previously described^44^. Temperature scans were collected with the following settings: 1□/min temperature gradient with a wait time of 30 s at the set temperature, then keeping the temperature within ± 0.1 of the target temperature for 10 s before starting the scan. Only one scan was acquired for all temperature ramp measurements. Samples of vesicles and protein were allowed to equilibrate for at least 15 minutes before spectra were acquired.

Equation 1 was used to determine the fraction of bound αSyn (w). Here, C_U_ is the average spectrum for unbound, disordered αSyn, C_S_ is the average saturated spectrum for membrane-bound αSyn, and C_I_ is the intermediate spectrum for αSyn that is partially bound to membranes (i.e., a mixture of bound and unbound states). Sonicated SUVs composed of 3:1 DPhPC/DPhPS were used to obtain saturated spectra, C_S_.

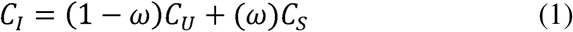

### Cryo-Electron Microscopy

For cryoEM grid preparation, 3 µl of purified sample was applied to freshly glow-discharged 1.2/1.3 µm holey carbon-coated 300-msh gold grids (Quantifoil), blotted for 2–4 s at 100% relative humidity and 4 °C, then vitrified in liquid ethane using a Vitrobot Mark IV (Thermo Fisher). Micrographs were collected with EPU data acquisition software (version 2.0) on a Glacios microscope (Thermo Fisher) operating at 200 keV. The microscope was equipped with a Falcon4 direct-electron detector. Images were acquired at 0.73 Å/ Å/pixel with a total electron dose of 50 electrons/Å^2 per image at 1.5 to 3.5 µm under focus.

### Curvatures from Cryo-EM of SUVs

To determine the curvature of the SUVs from cryoEM, the outer leaflet was manually traced using the Fiji plugin kappa and a B-spline fit was used to calculate curvature along the vesicle’s contour^70^. Applying the coordinates from kappa, we calculated the area (*A*) of the SUV using the Shoelace formula and determined the perimeter (*P*) by summing the Euclidean distance between adjacent coordinates. The circularity was defined as,

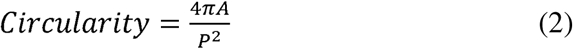

### Molecular Dynamics Simulations

To investigate the prevalence of membrane surface defects in gel-phase DPPC and liquid-phase DOPC membranes, as well as in mixtures doped with PS and PA, we constructed coarse-grained membrane systems using the Martini 3 force field parameters^71; 72^. These membranes were buckled to mimic the curvatures in the SUVs.

All Molecular Dynamics (MD) simulations were conducted using Gromacs 2025.2^73^. Symmetric bilayers were constructed with compositions according to Table 1^71;72^. To decrease computational time, an initial small system (8×8×25 nm) was created using Insane^74^. All systems were solvated with water and charged neutralized with. 15 M NaCl using Insane^74^. The initial small system was first minimized using a soft core steepest-descent minimization for 1000 steps with a 1.6 nm cutoff distance for short range neighbors, followed by an additional 1000 steps of conventional steepest descent minimization. To maintain bilayer structure, we performed successive steps of restrained equilibration where harmonic restraints were placed on the lipid headgroups to prevent lipid flip-flop. The time step was increased from 2-10 fs while restraints were weakened across the 55.25 ns of simulation. Following restrained equilibration, an additional 1 ns of unrestrained equilibration was performed with a timestep of 20 fs. We performed all simulations at 285K to replicate the phases of the membranes investigated in experiments (i.e., gel phase for DPPC and liquid for DOPC containing membranes)^75;76^. To generate the desired membrane size, we tessellated the small system 4-fold along the x-dimension, yielding a final system size of 32x8x25 nm, which we referred to as the large system. Final system compositions can be found in Table S1. The large systems were further minimized and equilibrated following the similar framework as the initial setup, but an additional 3 µs of unrestrained equilibration was performed with a timestep of 20 fs to generate the flat systems.

**Table 1:**
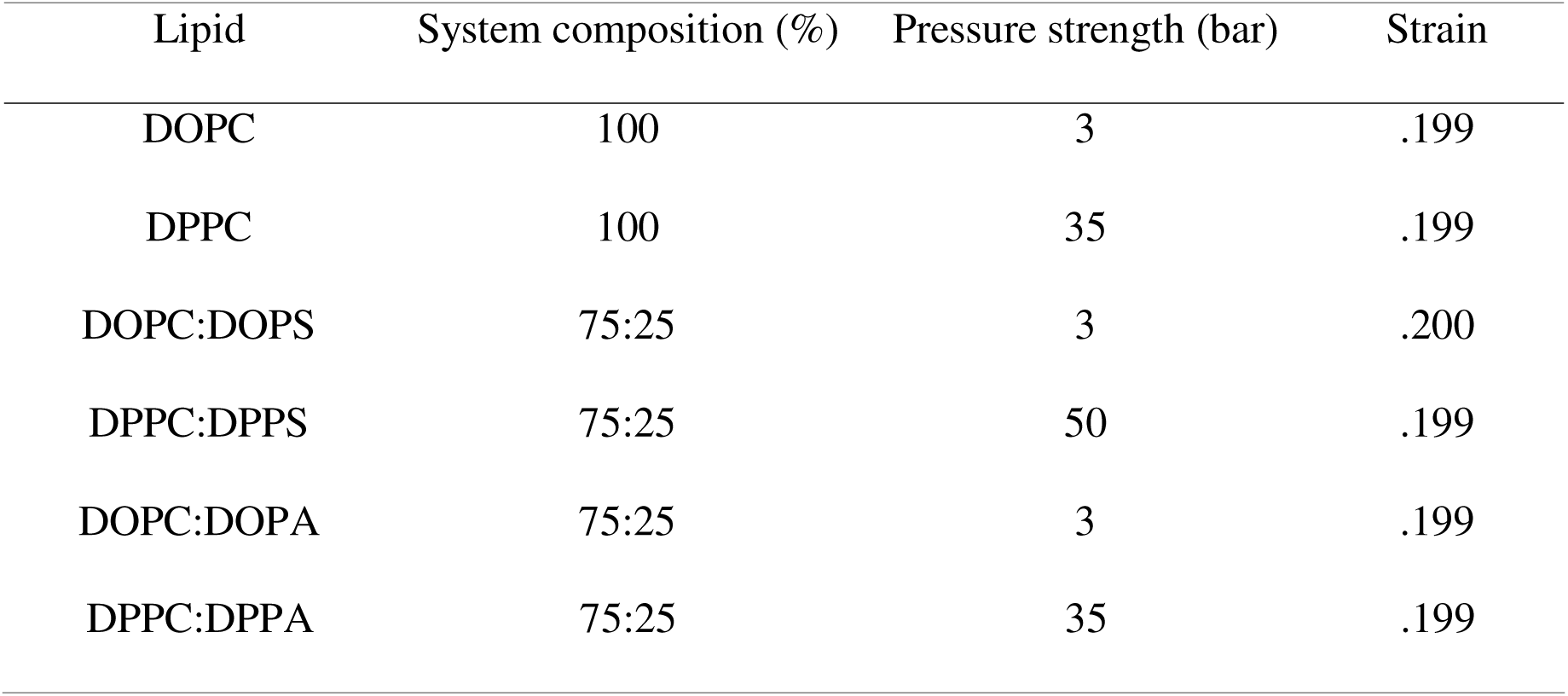
Table of lipid compositions.

To generate the buckled membranes, we compressed the flat system along the x-dimension using a Berendsen anisotropic barostat. Due to the mechanical properties of each membrane, the pressure required to induce reliable buckling depended on the lipid composition. Table S1 summarizes the pressure applied to induce buckling for each system. For all systems the buckling pressure was applied to the x-box dimension while the y-box dimension was held constant. The z-box dimension was coupled to a separate barostat with a compressibility of 3 × 10^−4^ bar^−1^ and pressure of 1 bar^77^.

To obtain the desired buckled membrane configurations, where curvatures were comparable between the different membrane compositions, a compression strain (*ɛ*) was calculated for each frame, where *L_0_* is the initial value of the box in the x-dimension and *L_t_* is the value of the box in the x-dimension at time, *t*.

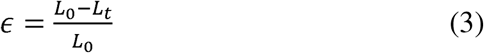

A compression strain of.2 was chosen for all buckled configurations to best match the SUV curvatures in EM. For each buckled system, the frame closest to a strain of .2 was selected as the starting configuration, and the exact strain values are listed in Table S1. *L_0_*was parameterized from the average box dimension of the corresponding flat membrane configurations during unrestrained equilibration in the NPT ensemble. In addition, this value was used to define the starting frame for the flat membrane configurations. Minimization, equilibration, and production runs were completed in the NVT ensemble for each system, to preserve the initial membrane geometry. A total of 1.8 μs of production for each buckled and flat system was completed and the final 600 ns was analyzed, as the defect surface coverage converged.

To ensure consistent alignment of the buckled region across trajectories, we centered the system by shifting the peak of the buckled membrane to the origin. We defined the peak as the set of lipids within 4 nm in height and within 1 nm in x-distance to the highest lipid. Restricting the x-dimension limited centering to a single peak. As PackMem previously established that defects persist for at most 20 ps, defects were computed for frames 100 ps apart to ensure defects in frames were independent and uncorrelated for later analysis^78^.

All codes for simulation set up are available on the GitHub (Table S2). The software versions used are documented in Table S2.

### Estimation of Curvature at the Peak of Buckled Membranes

To get an approximation of the curvatures of the buckled membranes, we identified the circle of best fit to the peak of the upper leaflet. We defined the peak by all lipids within 4 nm in height and ±4 nm in the x-dimension of the tallest lipid. For lipids in the peak region, we extracted the (x,y) coordinates of the PO4 beads and used the Pratt circle-fitting method to determine the circle of best fit, from which we calculated curvature^79^.

### Identification of Defects on Buckled and Flat Membranes

Since PackMem^78^ is restricted to flat membrane surfaces, we developed a workflow to relax this restriction using MDAnalysis^80;81^ and custom scripts. The software, termed Membrane Defect and Mapping Analysis (MemDMA), is available on GitHub (Table S2). In brief, we fit a quadrilateral mesh to the membrane’s surface and used its geometric information to analyze defects in curved and flat membrane geometries. To generate a representative surface of the membrane, the PO4 beads were selected and separated into leaflets according to Leafletfinder from MDAnalysis^80; 81^. A regular quadrilateral mesh with 1 Å spacing was fit to the point cloud, defined by the PO4 beads, using a piecewise cubic spline. To avoid handling defect detection across the periodic boundaries, we introduce a 10 Å buffer region which was excluded from the analysis. The normal vector for each quadrilateral was determined by averaging the normal vectors of the two triangles composing the quad. Lipids beads were projected along these normals onto the same plane as the quadrilateral to detect defects.

To determine which quads contained a defect, the algorithm was as follows: 1) identify the quad neighborhood for each bead 2) determine if the bead intersected the quad 3) classify if the quad was a defect depending on the identity of the beads mapped. To reduce computational cost, we restricted the set of quads onto which a given lipid could project. We started by selecting the closest quad to the PO4 bead of the lipid. For each bead of the current lipid, we projected the bead to any quad within 10 Å of the closest quad. The cutoff of 10 Å was chosen by comparing multiple cutoff values and selecting the smallest value such that no projections were missed. Beads located more than 6.36 Å from the quad were not projected. This depth cutoff was chosen to define shallow defects as those containing acyl chain beads within the plane of the polar beads, consistent with definitions used in PackMem^78^. The value of the cutoff corresponded to the thickness of the polar plane, defined as the sum of the GL2 diameter and PO4 radii. The radii for each bead were calculated by the *r_min_* from the Lennard-Jones potential, where σ was determined between two of the same beads.

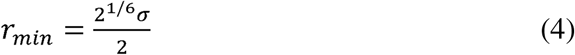

To determine if the bead overlapped with a quad we checked if the projected bead overlapped with any vertices or edges of the quad. If the bead overlapped with the quad, the quad’s value was incremented based on the bead type. As in PackMem, polar beads incremented the grid value by +1 while non-polar beads incremented the value by +.001. Previous work distinguishes between shallow and deep defects, where a quad value of 0 would define a deep defect while values between 0 and 1 as shallow defects^78^. However, this definition is dependent on an arbitrary choice of depth cut-off^78^. Therefore, we did not differentiate between shallow and deep defects during our defect analysis. Instead, a quad was assigned as a defect if the value was less than 1, consistent with other implementations of defect analysis tools^44; 76^.

### Defect Quantification and Statistics

Following the defect assignment, we applied the following filtering. Grids within ±8 Å of the membrane center along the y-dimension, along with all corresponding grids across the x-dimension, were removed from both flat and buckled membranes. This avoided the artifact where a polar bead always occupied the central grid due to our buckled membrane centering strategy. Following PackMem, we excluded defects with an area less than 15 Å^2^ from the analysis, as small defects are similarly distributed between systems^78^. We performed all analyses on the upper leaflet, as it corresponds to the experimentally relevant curvatures.

We calculated defect surface coverage and average defect size for the entire membrane and the peak region of the membrane. The entire membrane was defined as all membrane grids, while the peak region comprised those within ±2.5 nm of the membrane’s center along the x-dimension. Defect surface coverage was calculated by dividing the number of grids containing a defect by the total number of grids in the defined membrane region. The average defect size was obtained by first computing the mean defect size per frame and then averaging these values across all frames in the trajectory. To characterize the probability distribution of defects sizes, we calculated the log probability of observing a defect of a given size following a framework similar to PackMem^78^. Defect sizes were binned by 1 Å² intervals, normalized by the total number of defects in the trajectory, and the logarithm of the resulting probabilities was taken.

Given that the membranes are buckled along the x-direction, we evaluated the defect propensity as a function of x. Since the geometries of the membrane strips are small in the y-dimension, we assume that the membranes are approximately planar in the y-direction. We therefore compute the defect likelihood as a function of x-position averaged over samples in y and over all frames. As the membrane’s height increased with curvature, the average membrane height along the x-dimension was used as a proxy for curvature.

### FRAP Analysis

Equation 5 was used to determine *x_m,_*the fraction of mobile proteins, and *k_off_*, the rate of αSyn unbinding. Here, B is the average fluorescence intensity of the ROI, B_∞_ is the average intensity of the ROI at steady-state recovery, *t* is time, and *k_off_*represents the off-rate of the mobile fraction, and *k_off,i_* represents the off-rate of the immobile fraction of protein coming off the membrane.

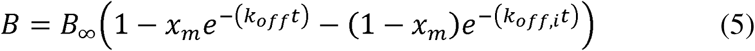

The normalized initial rate was defined as the product of *x_m_* and *k_off_*.

### Fluorescence Anisotropy

Fluorescence anisotropy data were acquired using a JASCO FP-8500 spectrofluorometer. SUVs containing DPPC (60 μM), 3:1 DPPC:DPPS (60 μM), and 3:1 DPPC:DPPA (60 μM) were prepared by extrusion with 50 nm membranes as described above (under “SUV preparation”). DPPC and DPPC:DPPS extrusions were carried out at 60□, and DPPC:DPPA extrusions were carried out at 70□ to ensure SUV formation above the melting temperature. The average diameter of vesicles used for fluorescence anisotropy across all compositions was 69.8 ± 4.2 nm, determined by dynamic light scattering. A stock DPH solution (200 μm) was added to the SUVs in sodium phosphate buffer (20 mM sodium phosphate, 150 mM NaCl, pH 7.0) at phospholipid:DPH molar ratios of 180:1. A spectrofluorometer (JASCO FP-8500) was used to excite DPH at 355 nm, and its fluorescence emission intensity at 425 nm and anisotropy value were monitored as the temperature varied from 20 to 70□. A 250:1 lipid to protein ratio was used for experiments containing αSyn.

Equation 6 was used to fit the anisotropy data and determine the melting temperature, *T_m_*.

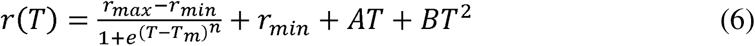

Here, *r* is the anisotropy value, *T* is the temperature, *n* is the Hill coefficient, and *A* and *B* are regression parameters for baseline correction.

## Acknowledgments

Thank you to Htet A. Khant and the Core Center of Excellence in Nano Imaging at USC for assistance with cryoEM imaging. Thank you to Jennifer C. Lee for generously providing the *pRK172 human* α*Syn(Y136C)* plasmid. Thank you to Peter J. Chung for providing the pNatB plasmid. This research was supported by the National Institutes of Health through R35GM147333 to O.H.K., D.H.J., and W.F.Z. C.M.S is supported in part by the Pathways in Biological Science Training Program, National Institutes of Health T32-GM133351. Molecular dynamics simulations were run on hardware hosted by the Triton Shared Computing Cluster. We are also grateful to the UCSD Physics Computing Facility for computational resources.

## References

1. Maroteaux L, Campanelli J, Scheller R. 1988. Synuclein: A neuron-specific protein localized to the nucleus and presynaptic nerve terminal. Journal of Neuroscience. 8(8).

2. Venda LL, Cragg SJ, Buchman VL, Wade-Martins R. 2010. Α-synuclein and dopamine at the crossroads of parkinson’s disease. Trends in Neurosciences. 33(12).

3. Galvagnion C. 2017. The role of lipids interacting with α-synuclein in the pathogenesis of parkinson’s disease. Journal of Parkinson’s Disease. 7(3).

4. Spillantini MG, Schmidt ML, Lee VM-Y, Trojanowski JQ, Jakes R, Goedert M. 1997. Α-synuclein in lewy bodies. Nature. 388(6645):839–840.

5. Spillantini MG, Crowther RA, Jakes R, Hasegawa M, Goedert M. 1998. Α-synuclein in filamentous inclusions of lewy bodies from parkinson’s disease and dementia with lewy bodies. Proceedings of the National Academy of Sciences. 95(11):6469–6473.

6. Nakamura K, Nemani VM, Wallender EK, Kaehlcke K, Ott M, Edwards RH. 2008. Optical reporters for the conformation of α-synuclein reveal a specific interaction with mitochondria. Journal of Neuroscience. 28(47).

7. Burré J. 2015. The synaptic function of α-synuclein. Journal of Parkinson’s Disease. 5(4).

8. Kaur U, Lee JC. 2020. Unroofing site-specific α-synuclein–lipid interactions at the plasma membrane. Proceedings of the National Academy of Sciences. 117(32):18977–18983.

9. Sarchione A, Marchand A, Taymans J-M, Chartier-Harlin M-C. 2021. Alpha-synuclein and lipids: The elephant in the room? Cells 2021, Vol 10,. 10(9).

10. Barbuti PA, Guardia-Laguarta C, Yun T, Chatila ZK, Flowers X, Wong C, Santos BFR, Larsen SB, Lotti JS, Hattori N et al. 2025. The role of alpha-synuclein in synucleinopathy: Impact on lipid regulation at mitochondria–er membranes. npj Parkinson’s Disease. 11(1).

11. Burré J, Sharma M, Tsetsenis T, Buchman V, Etherton MR, Südhof TC. 2010. Α-synuclein promotes snare-complex assembly in vivo and in vitro. Science. 329(5999):1663–1667.

12. Snead D, Eliezer D. 2014. Alpha-synuclein function and dysfunction on cellular membranes. Experimental Neurobiology. 23(4):292–313.

13. Galvagnion C, Buell AK, Meisl G, Michaels TCT, Vendruscolo M, Knowles TPJ, Dobson CM. 2015. Lipid vesicles trigger α-synuclein aggregation by stimulating primary nucleation. Nature Chemical Biology. 11(3):229–234.

14. Khounlo R, Hawk BJD, Khu T-M, Yoo G, Lee NK, Pierson J, Shin Y-K. 2021. Membrane binding of α-synuclein stimulates expansion of snare-dependent fusion pore. Frontiers in Cell and Developmental Biology. 9.

15. Man WK, Tahirbegi B, Vrettas MD, Preet S, Ying L, Vendruscolo M, De Simone A, Fusco G. 2021. The docking of synaptic vesicles on the presynaptic membrane induced by α-synuclein is modulated by lipid composition. Nature Communications. 12(1).

16. Sharma M, Burré J. 2023. Α-synuclein in synaptic function and dysfunction. Trends in Neurosciences. 46(2).

17. Davidson WS, Jonas A, Clayton DF, George JM. 1998. Stabilization of α-synuclein secondary structure upon binding to synthetic membranes. Journal of Biological Chemistry. 273(16):9443–9449.

18. Chandra S, Chen X, Rizo J, Jahn R, Südhof TC. 2003. A broken α-helix in folded α-synuclein. Journal of Biological Chemistry. 278(17):15313–15318.

19. Middleton ER, Rhoades E. 2010. Effects of curvature and composition on α-synuclein binding to lipid vesicles. Biophysical Journal. 99(7):2279–2288.

20. Stöckl M, Fischer P, Wanker E, Herrmann A. 2008. Α-synuclein selectively binds to anionic phospholipids embedded in liquid-disordered domains. Journal of Molecular Biology. 375(5).

21. Hellstrand E, Grey M, Ainalem M-L, Ankner J, Forsyth VT, Fragneto G, Haertlein M, Dauvergne M-T, Nilsson H, Brundin P et al. 2013. Adsorption of α-synuclein to supported lipid bilayers: Positioning and role of electrostatics.

22. Andersson A, Linse S, Sparr E, Fornasier M, Jönsson P. 2024. The density of anionic lipids modulates the adsorption of α-synuclein onto lipid membranes. Biophysical Chemistry. 305.

23. Pandey AP, Haque F, Rochet J-C, Hovis JS. 2009. Clustering of α-synuclein on supported lipid bilayers: Role of anionic lipid, protein, and divalent ion concentration. Biophysical Journal. 96(2):540–551.

24. O’Leary EI, Jiang Z, Strub M-P, Lee JC. 2018. Effects of phosphatidylcholine membrane fluidity on the conformation and aggregation of n-terminally acetylated α-synuclein. Journal of Biological Chemistry. 293(28):11195–11205.

25. Bartels T, Ahlstrom LS, Leftin A, Kamp F, Haass C, Brown MF, Beyer K. 2010. The n-terminus of the intrinsically disordered protein α-synuclein triggers membrane binding and helix folding. Biophysical Journal. 99(7):2116–2124.

26. Kjaer L, Giehm L, Heimburg T, Otzen D. 2009. The influence of vesicle size and composition on α-synuclein structure and stability. Biophysical Journal. 96(7):2857–2870.

27. Kamp F, Beyer K. 2006. Binding of α-synuclein affects the lipid packing in bilayers of small vesicles. Journal of Biological Chemistry. 281(14):9251–9259.

28. Nuscher B, Kamp F, Mehnert T, Odoy S, Haass C, Kahle PJ, Beyer K. 2004. Α-synuclein has a high affinity for packing defects in a bilayer membrane. Journal of Biological Chemistry. 279(21):21966–21975.

29. Shvadchak VV, Falomir-Lockhart LJ, Yushchenko DA, Jovin TM. 2011. Specificity and kinetics of α-synuclein binding to model membranes determined with fluorescent excited state intramolecular proton transfer (esipt) probe. Journal of Biological Chemistry. 286(15):13023–13032.

30. Levi M, Wilson P, Nguyen S, Iorio E, Sapora O, Parasassi T. 1997. In k562 and hl60 cells membrane ageing during cell growth is associated with changes in cholesterol concentration. Mechanisms of Ageing and Development. 97(2).

31. Giusto NM, Salvador GA, Castagnet PI, Pasquaré SJ, Ilincheta de Boschero MG, Giusto NM, Salvador GA, Castagnet PI, Pasquaré SJ, Ilincheta de Boschero MG. 2002. Age-associated changes in central nervous system glycerolipid composition and metabolism. Neurochemical Research 2002 27:11. 27(11).

32. Fabelo N, Martín V, Santpere G, Marín R, Torrent L, Ferrer I, Díaz M. 2011. Severe alterations in lipid composition of frontal cortex lipid rafts from parkinson’s disease and incidental parkinson’s disease. Molecular Medicine. 17(9-10):1107–1118.

33. Pranke IM, Morello V, Bigay J, Gibson K, Verbavatz J-M, Antonny B, Jackson CL. 2011. Α-synuclein and alps motifs are membrane curvature sensors whose contrasting chemistry mediates selective vesicle binding. Journal of Cell Biology. 194(1):89–103.

34. Johnson DH, Kou OH, Bouzos N, Zeno WF. 2024. Protein–membrane interactions: Sensing and generating curvature. Trends in Biochemical Sciences. 49(5).

35. Dikiy I, Eliezer D. 2014. N-terminal acetylation stabilizes n-terminal helicity in lipid– and micelle-bound α-synuclein and increases its affinity for physiological membranes. Journal of Biological Chemistry. 289(6):3652–3665.

36. Maltsev AS, Ying J, Bax A. 2012. Impact of n-terminal acetylation of α-synuclein on its random coil and lipid binding properties.

37. Burré J, Vivona S, Diao J, Sharma M, Brunger AT, Südhof TC. 2013. Properties of native brain α-synuclein. Nature. 498(7453):E4–E6.

38. Theillet F-X, Binolfi A, Bekei B, Martorana A, Rose HM, Stuiver M, Verzini S, Lorenz D, Van Rossum M, Goldfarb D, et al. 2016. Structural disorder of monomeric α-synuclein persists in mammalian cells. Nature. 530(7588):45–50.

39. Arnesen T, Van Damme P, Polevoda B, Helsens K, Evjenth R, Colaert N, Varhaug JE, Vandekerckhove J, Lillehaug JR, Sherman F et al. 2009. Proteomics analyses reveal the evolutionary conservation and divergence of n-terminal acetyltransferases from yeast and humans. Proceedings of the National Academy of Sciences. 106(20):8157–8162.

40. Anderson JP, Walker DE, Goldstein JM, De Laat R, Banducci K, Caccavello RJ, Barbour R, Huang J, Kling K, Lee M et al. 2006. Phosphorylation of ser-129 is the dominant pathological modification of α-synuclein in familial and sporadic lewy body disease. Journal of Biological Chemistry. 281(40):29739–29752.

41. Öhrfelt A, Zetterberg H, Andersson K, Persson R, Secic D, Brinkmalm G, Wallin A, Mulugeta E, Francis PT, Vanmechelen E et al. 2011. Identification of novel α-synuclein isoforms in human brain tissue by using an online nanolc-esi-fticr-ms method. Neurochemical Research. 36(11):2029–2042.

42. Runfola M, De Simone A, Vendruscolo M, Dobson CM, Fusco G. 2020. The n-terminal acetylation of α-synuclein changes the affinity for lipid membranes but not the structural properties of the bound state. Scientific Reports. 10(1).

43. Hatzakis NS, Bhatia VK, Larsen J, Madsen KL, Bolinger P-Y, Kunding AH, Castillo J, Gether U, Hedegård P, Stamou D et al. 2009. How curved membranes recruit amphipathic helices and protein anchoring motifs. Nature Chemical Biology 2009 5:11. 5(11).

44. Johnson DH, Kou OH, White JM, Ramirez SY, Margaritakis A, Chung PJ, Jaeger VW, Zeno WF. 2025. Lipid packing defects are necessary and sufficient for membrane binding of α-synuclein. Communications Biology. 8(1).

45. Zeno WF, Day KJ, Gordon VD, Stachowiak JC. 2020. Principles and applications of biological membrane organization. Annual Review of Biophysics. 49(Volume 49, 2020).

46. Hirst LS, Ossowski A, Fraser M, Geng J, Selinger JV, Selinger RLB. 2013. Morphology transition in lipid vesicles due to in-plane order and topological defects. Proceedings of the National Academy of Sciences. 110(9):3242–3247.

47. Ickenstein LM, Arfvidsson MC, Needham D, Mayer LD, Edwards K. 2003. Disc formation in cholesterol-free liposomes during phase transition. Biochimica et Biophysica Acta (BBA) – Biomembranes. 1614(2).

48. Andersson M, Hammarstroem L, Edwards K. 1995. Effect of bilayer phase transitions on vesicle structure, and its influence on the kinetics of viologen reduction.

49. Van Rooijen BD, Claessens MMAE, Subramaniam V. 2009. Lipid bilayer disruption by oligomeric α-synuclein depends on bilayer charge and accessibility of the hydrophobic core. Biochimica et Biophysica Acta (BBA) – Biomembranes. 1788(6).

50. Jo E, Mclaurin J, Yip CM, St. George-Hyslop P, Fraser PE. 2000. Α-synuclein membrane interactions and lipid specificity. Journal of Biological Chemistry. 275(44):34328–34334.

51. Van Rooijen BD, Claessens MMAE, Subramaniam V. 2008. Membrane binding of oligomeric α synuclein depends on bilayer charge and packing. FEBS Letters. 582(27):3788–3792.

52. Sonntag Y, Musgaard M, Olesen C, Schiøtt B, Møller JV, Nissen P, Thøgersen L. 2011. Mutual adaptation of a membrane protein and its lipid bilayer during conformational changes. Nature Communications. 2(1):304.

53. Bernhardt N, Ozturk TN, Zhang S, Schwartz N, Chadda R, Gil-Ley A, Robertson JL, Faraldo-GóMez JD. 2025. Molecular basis for the regulation of membrane proteins through preferential lipid solvation. Nature Chemical Biology.

54. Levental I, Lyman E. 2023. Regulation of membrane protein structure and function by their lipid nano-environment. Nature Reviews Molecular Cell Biology. 24(2):107–122.

55. Sych T, Levental KR, Sezgin E. 2022. Lipid–protein interactions in plasma membrane organization and function. Annual Review of Biophysics. 51(1):135–156.

56. Garten M, Prévost C, Cadart C, Gautier R, Bousset L, Melki R, Bassereau P, Vanni S. 2015. Methyl-branched lipids promote the membrane adsorption of α-synuclein by enhancing shallow lipid-packing defects. Physical Chemistry Chemical Physics. 17(24):15589–15597.

57. Ouberai MM, Wang J, Swann MJ, Galvagnion C, Guilliams T, Dobson CM, Welland ME. 2013. Α-synuclein senses lipid packing defects and induces lateral expansion of lipids leading to membrane remodeling. Journal of Biological Chemistry. 288(29):20883–20895.

58. Nagarajan S, Schuler EE, Ma K, Kindt JT, Dyer RB. 2012. Dynamics of the gel to fluid phase transformation in unilamellar dppc vesicles.

59. Kranenburg M, Smit B. 2005. Phase behavior of model lipid bilayers. The Journal of Physical Chemistry B. 109(14).

60. Meyer HW. 1996. Pretransition-ripples in bilayers of dipalmitoylphosphatidylcholine: Undulation or periodic segments? A freeze-fracture study. Biochimica et Biophysica Acta (BBA) – Lipids and Lipid Metabolism. 1302(2).

61. Cunningham BA, Brown A-D, Wolfe DH, Williams WP, Brain A. 1998. Ripple phase formation in phosphatidylcholine: Effect of acyl chain relative length, position, and unsaturation. Physical Review E. 58(3).

62. Kaasgaard T, Leidy C, Crowe JH, Mouritsen OG, Jørgensen K. 2003. Temperature-controlled structure and kinetics of ripple phases in one– and two-component supported lipid bilayers. Biophysical Journal. 85(1):350–360.

63. Foley SL, Hossein A, Deserno M. 2022. Fluid-gel coexistence in lipid membranes under differential stress. Biophysical Journal. 121(16).

64. Rappolt M, Pabst G, Rapp G, Kriechbaum M, Amenitsch H, Krenn C, Bernstorff S, Laggner P. 2000. New evidence for gel-liquid crystalline phase coexistence in the ripple phase of phosphatidylcholines. European Biophysics Journal 2000 29:2. 29(2).

65. Momin N, Lee S, Gadok AK, Busch DJ, Bachand GD, Hayden CC, Stachowiak JC, Sasaki DY. 2015. Designing lipids for selective partitioning into liquid ordered membrane domains. Soft Matter. 11(16):3241–3250.

66. Johnson M, Coulton AT, Geeves MA, Mulvihill DP. 2010. Targeted amino-terminal acetylation of recombinant proteins in e. Coli. PLoS ONE. 5(12):e15801.

67. Zeno WF, Baul U, Snead WT, Degroot ACM, Wang L, Lafer EM, Thirumalai D, Stachowiak JC. 2018. Synergy between intrinsically disordered domains and structured proteins amplifies membrane curvature sensing. Nature Communications. 9(1).

68. Zeno WF, Thatte AS, Wang L, Snead WT, Lafer EM, Stachowiak JC. 2019. Molecular mechanisms of membrane curvature sensing by a disordered protein. Journal of the American Chemical Society. 141(26).

69. Aguet F, Antonescu N, Costin, Mettlen M, Schmid L, Sandra, Danuser G. 2013. Advances in analysis of low signal-to-noise images link dynamin and ap2 to the functions of an endocytic checkpoint. Developmental Cell. 26(3):279–291.

70. Kappa. 2013. [accessed 2026]. https://github.com/fiji/Kappa.

71. Souza PCT, Alessandri R, Barnoud J, Thallmair S, Faustino I, Grünewald F, Patmanidis I, Abdizadeh H, Bruininks BMH, Wassenaar TA et al. 2021. Martini 3: A general purpose force field for coarse-grained molecular dynamics. Nature Methods. 18(4):382–388.

72. Pedersen KB, Ingólfsson HI, Ramirez-Echemendia DP, Borges-Araújo L, Andreasen MD, Empereur-mot C, Melcr J, Ozturk TN, Bennett WFD, Kjølbye LR et al. 2025. The martini 3 lipidome: Expanded and refined parameters improve lipid phase behavior. ACS Central Science. 11(9).

73. Gromacs 2025.2 manual. 2025. [accessed 2026]. https://zenodo.org/records/15387070.

74. Wassenaar TA, Ingólfsson HI, Böckmann RA, Tieleman DP, Marrink SJ. 2015. Computational lipidomics with insane: A versatile tool for generating custom membranes for molecular simulations. Journal of Chemical Theory and Computation. 11(5).

75. Jaschonek S, Cascella M, Gauss J, Diezemann G, Milano G. 2018. Intramolecular structural parameters are key modulators of the gel-liquid transition in coarse grained simulations of dppc and dopc lipid bilayers. Biochemical and Biophysical Research Communications. 498(2).

76. Van Der Pol RWI, Brinkmann BW, Sevink GJA. 2024. Analyzing lipid membrane defects via a coarse-grained to triangulated surface map: The role of lipid order and local curvature in molecular binding. Journal of Chemical Theory and Computation. 20(7):2888–2900.

77. Stroh KS, Risselada HJ. 2021. Quantifying membrane curvature sensing of peripheral proteins by simulated buckling and umbrella sampling. Journal of Chemical Theory and Computation. 17(8).

78. Gautier R, Bacle A, Tiberti ML, Fuchs PF, Vanni S, Antonny B. 2018. Packmem: A versatile tool to compute and visualize interfacial packing defects in lipid bilayers. Biophysical Journal. 115(3):436–444.

79. Circle fit (pratt method). 2026. MATLAB Central File Exchange; [accessed 2026]. https://www.mathworks.com/matlabcentral/fileexchange/22643-circle-fit-pratt-method.

80. Gowers RJ, Linke M, Barnoud J, Reddy TJE, Melo MN, Seyler SL, Domański J, Dotson DL, Buchoux S, Kenney IM, et al. 2016. Mdanalysis: A python package for the rapid analysis of molecular dynamics simulations. SciPy.

81. Michaud Agrawal N, Denning EJ, Woolf TB, Beckstein O. 2011. Mdanalysis: A toolkit for the analysis of molecular dynamics simulations. Journal of Computational Chemistry. 32(10):2319–2327.

