## Supplemental Figures for "Membrane Phase, Charge, and Curvature Regulate α-Synuclein Binding Dynamics"

**Supplementary Information for**
Membrane Phase and Charge Regulate α-Synuclein Binding and Curvature Sensing

Orianna H. Kou, Cailyn M. Sakurai, Stephanie Y. Ramirez, Brian H. Kim, David H. Johnson, Zixin Zhang, Christopher T. Lee, Wade F. Zeno*

*Corresponding Author: Wade F. Zeno


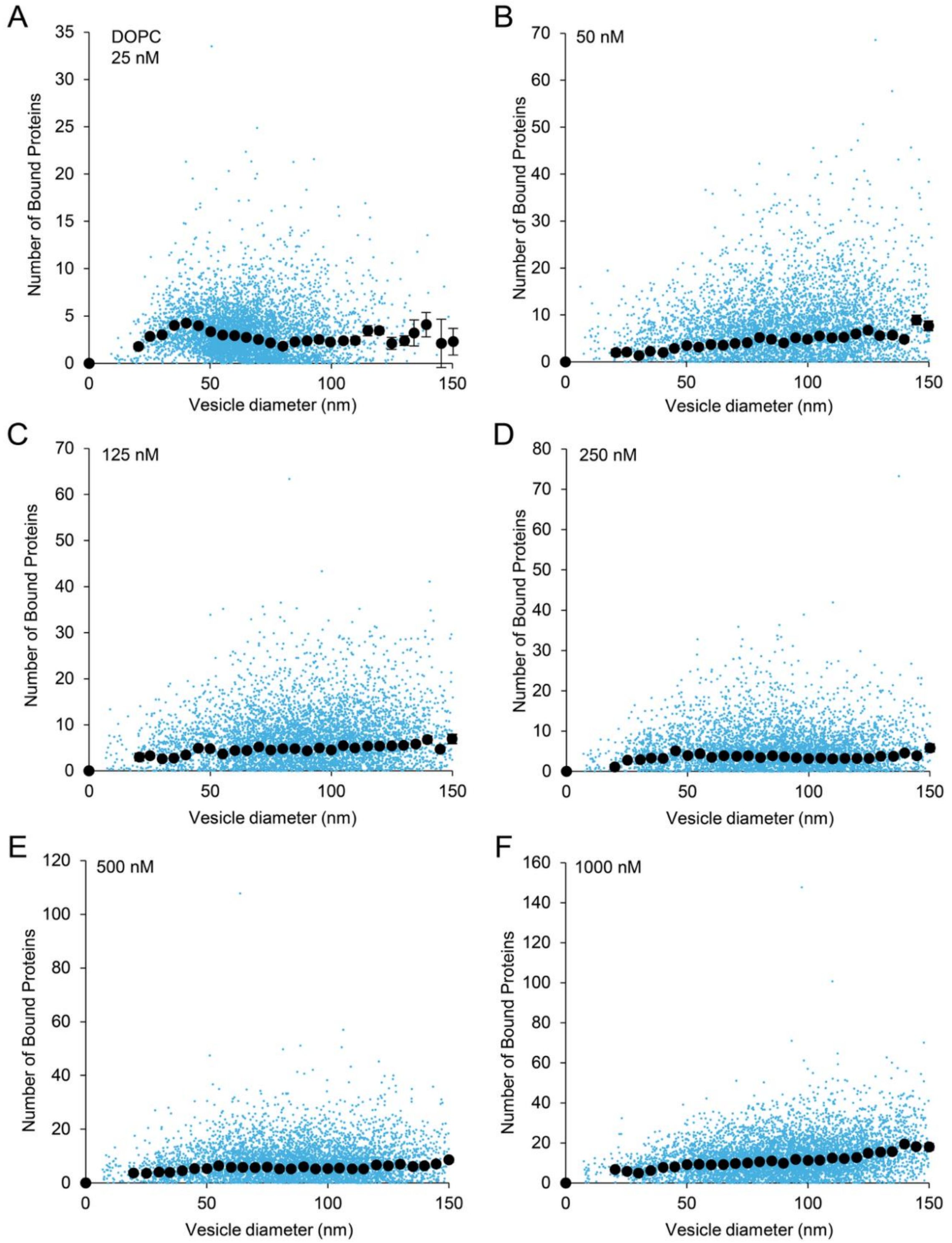


**Figure S1. Raw data for protein binding on DOPC vesicles.** The distribution of proteins bound to individual DOPC vesicles (blue circles) and the average number of proteins bound in various size bins (black circles) at (A) 25 nM, (B) 50 nM, (C) 125 nM, (D) 250 nM, (E) 500 nM, and (F) 1000 nM. Error bars represent the standard error of the mean.

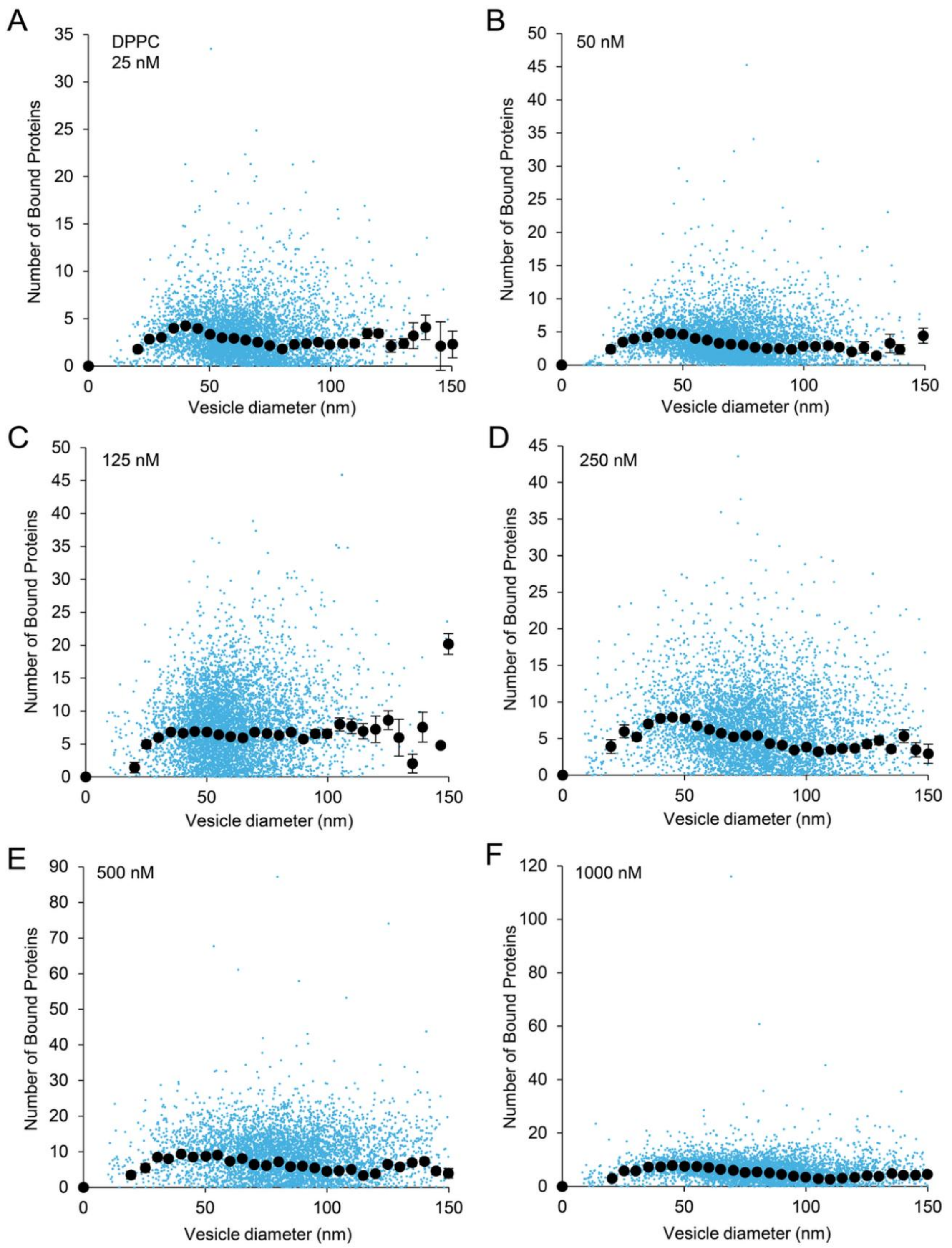


**Figure S2. Raw data for protein binding on DPPC vesicles.** The distribution of proteins bound to individual DPPC vesicles (blue circles) and the average number of proteins bound in various size bins (black circles) at (A) 25 nM, (B) 50 nM, (C) 125 nM, (D) 250 nM, (E) 500 nM, and (F) 1000 nM. Error bars represent the standard error of the mean.

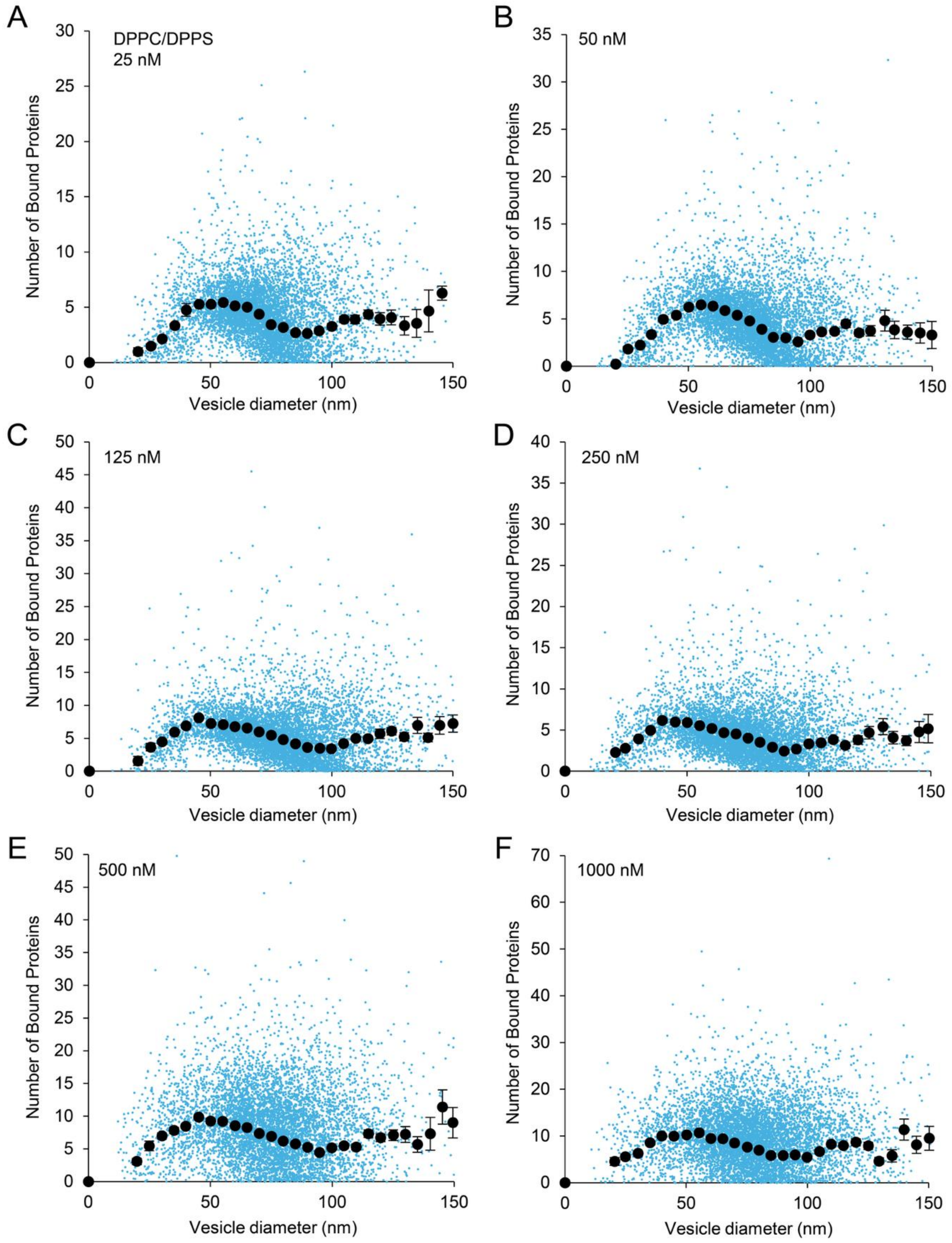


**Figure S3.** Raw data for protein binding on DPPC/DPPS vesicles**.** The distribution of proteins bound to individual 3:1 DPPC:DPPS vesicles (blue circles) and the average number of proteins bound in various size bins (black circles) at (A) 25 nM, (B) 50 nM, (C) 125 nM, (D) 250 nM, (E) 500 nM, and (F) 1000 nM. Error bars represent the standard error of the mean.

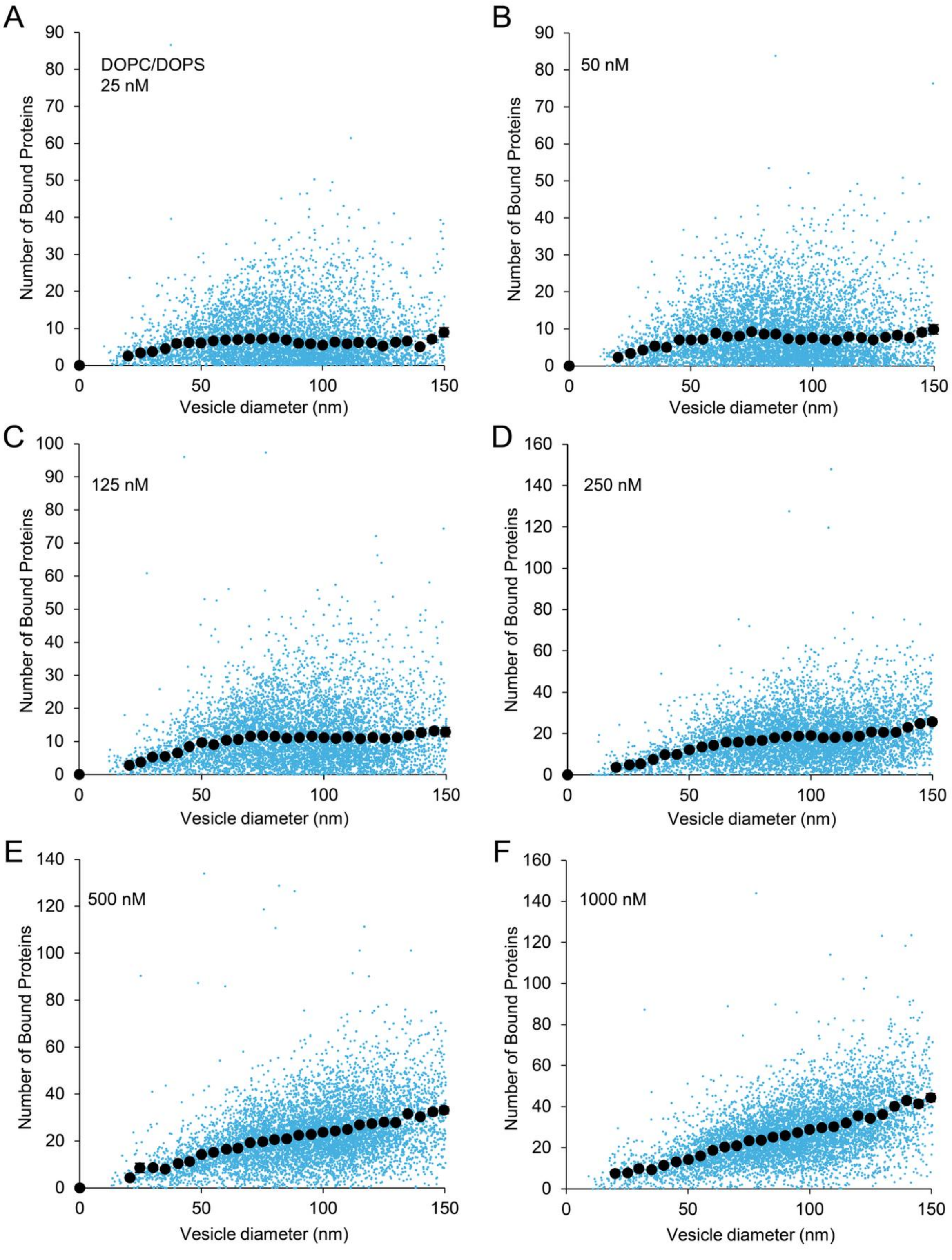


**Figure S4. Raw data for protein binding on DOPC/DOPS vesicles.** The distribution of proteins bound to individual 3:1 DOPC:DOPS vesicles (blue circles) and the average number of proteins bound in various size bins (black circles) at (A) 25 nM, (B) 50 nM, (C) 125 nM, (D) 250 nM, (E) 500 nM, and (F) 1000 nM. Error bars represent the standard error of the mean.

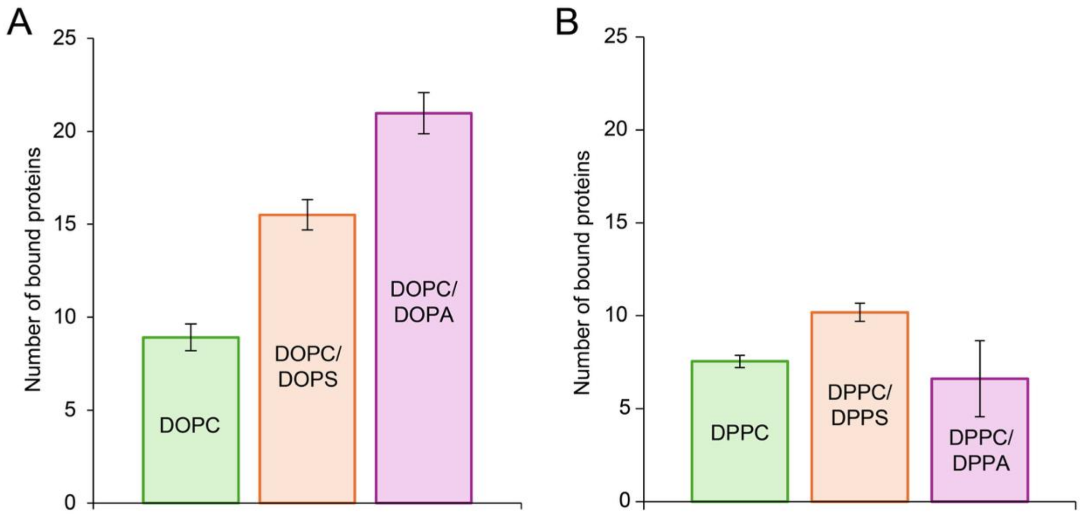


**Figure S5. Fluorescence microscopy reveals that PA lipids enhance αSyn binding to liquid-phase membranes, but not gel-phase membranes.** (A-B) Association of 1μM αSyn to SUVs with an average diameter of 50 nm. Error bars represent the 99% confidence interval of the mean.

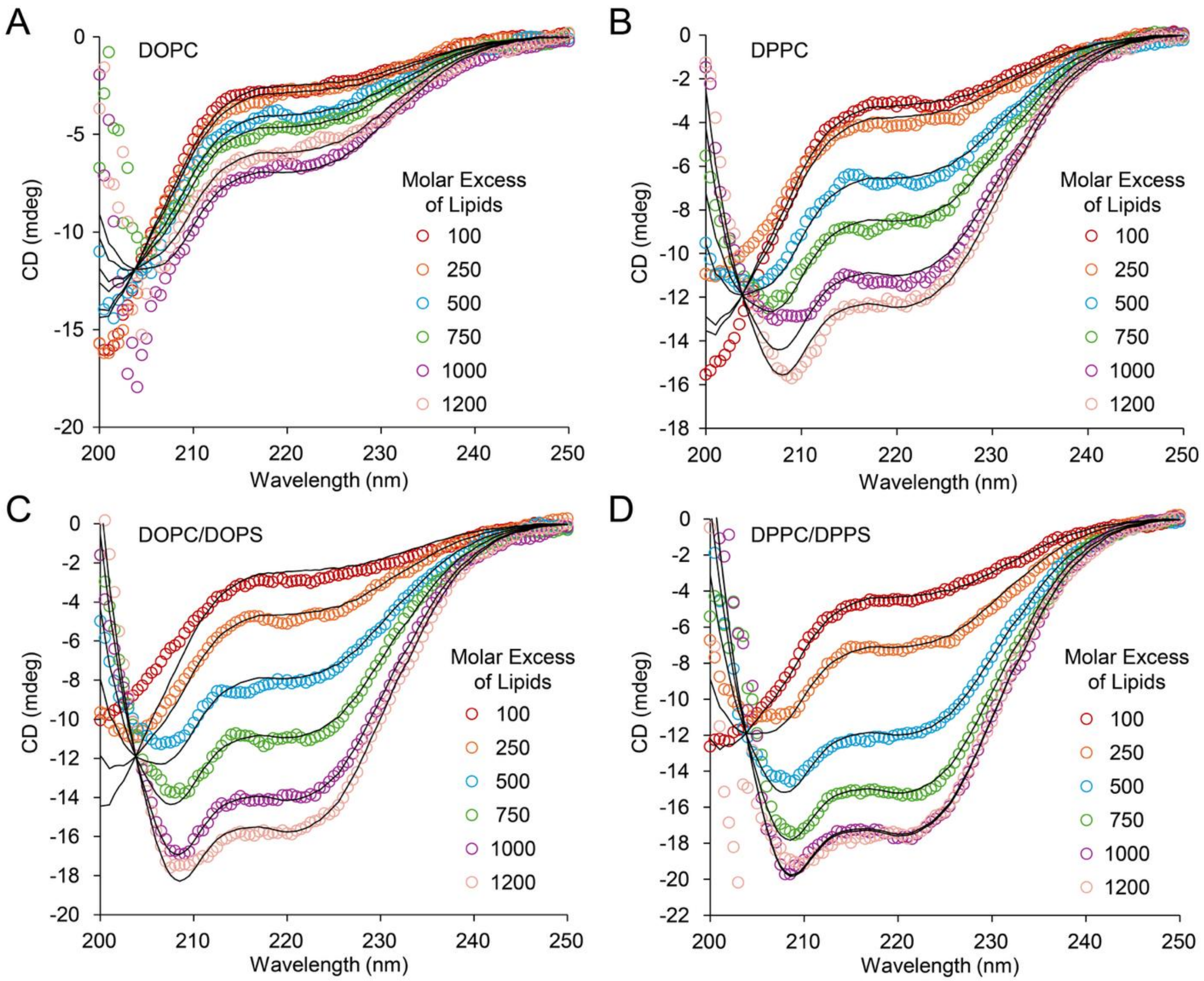


**Figure S6.** Circular dichroism spectra for αSyn bound to vesicles composed of (A) DOPC, (B) DPPC, (C) 3:1 DPPC/DPPS, and (D) 3:1 DOPC/DOPS. Molar excess of lipid is the lipid molar concentration divided by protein molar concentration. 5 μM of protein was used for these experiments.

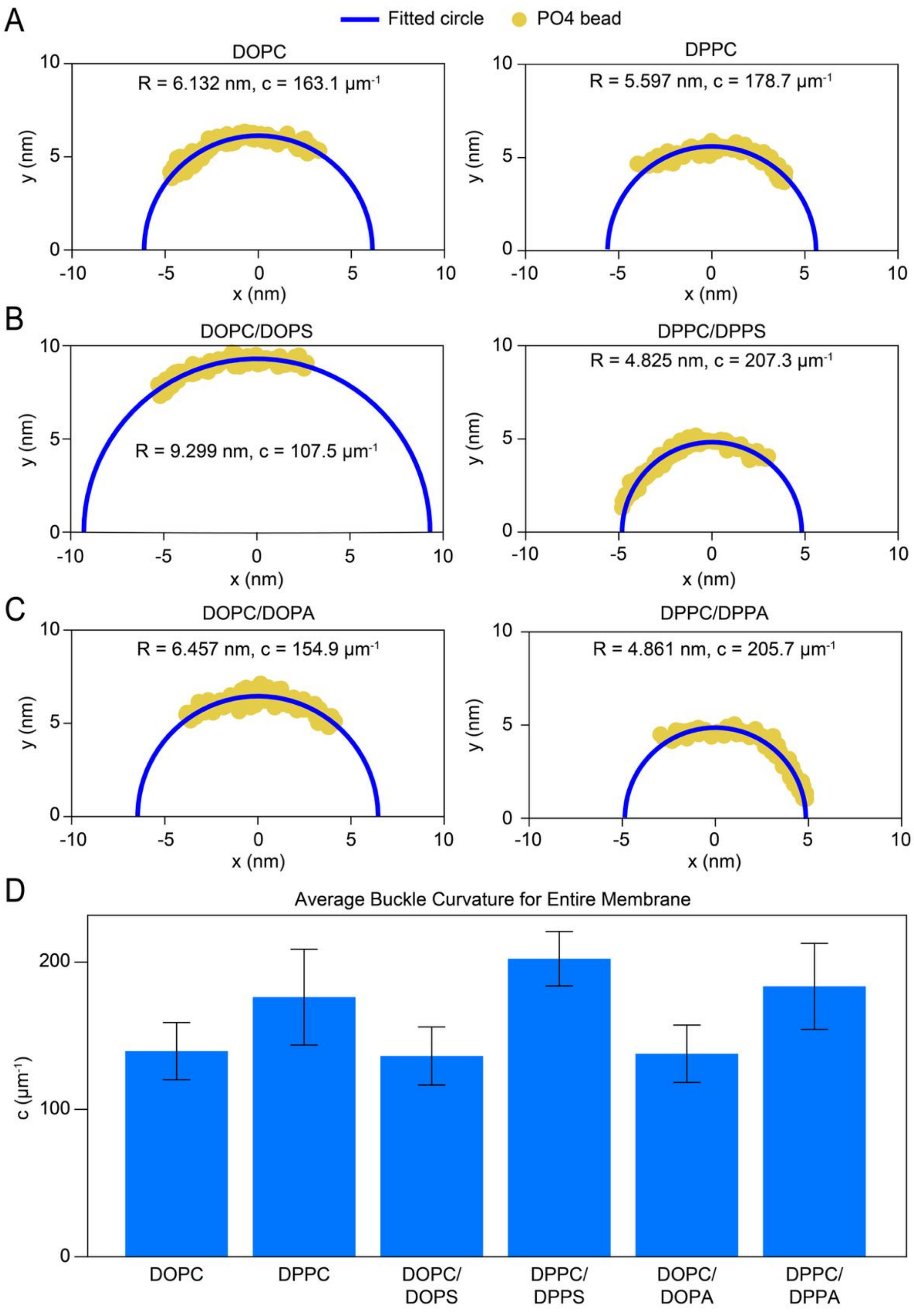


**Figure S7.** (A-C) Representative curvatures of the various simulated liquid- and gel-phase compositions, which directly correspond to the representative MD images in Figure 3 and Supplementary Figure 10-11, determined using Pratt circle-fitting. (D) Average curvature for the 6 simulated compositions. Error bars represent the standard deviations calculated over all frames of the simulation.

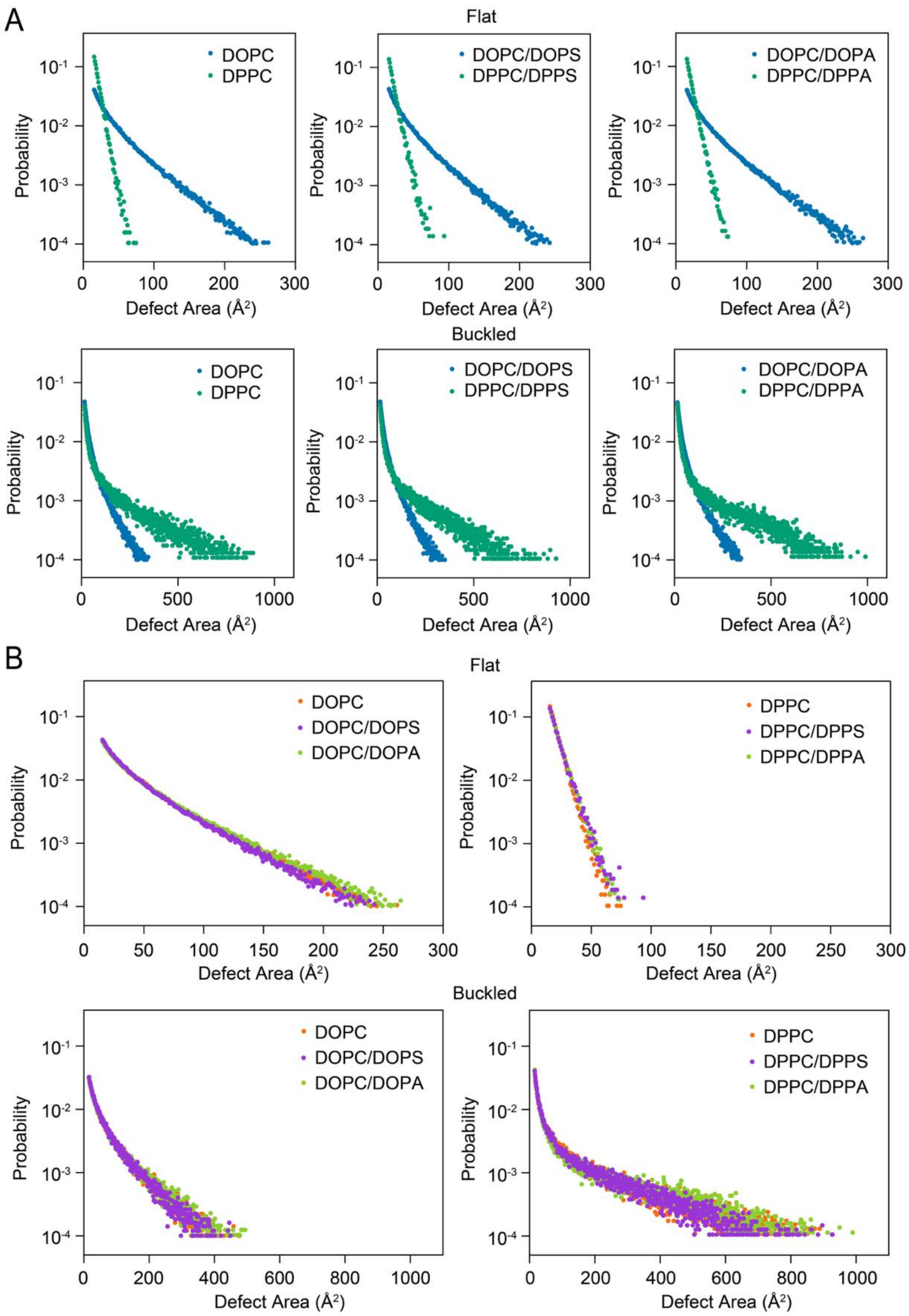


**Figure S8.** Comparison of defect area probability distribution across the entire membrane (A) between phases and (B) within each phase as a function of headgroup and membrane curvature (i.e., flat or buckled).

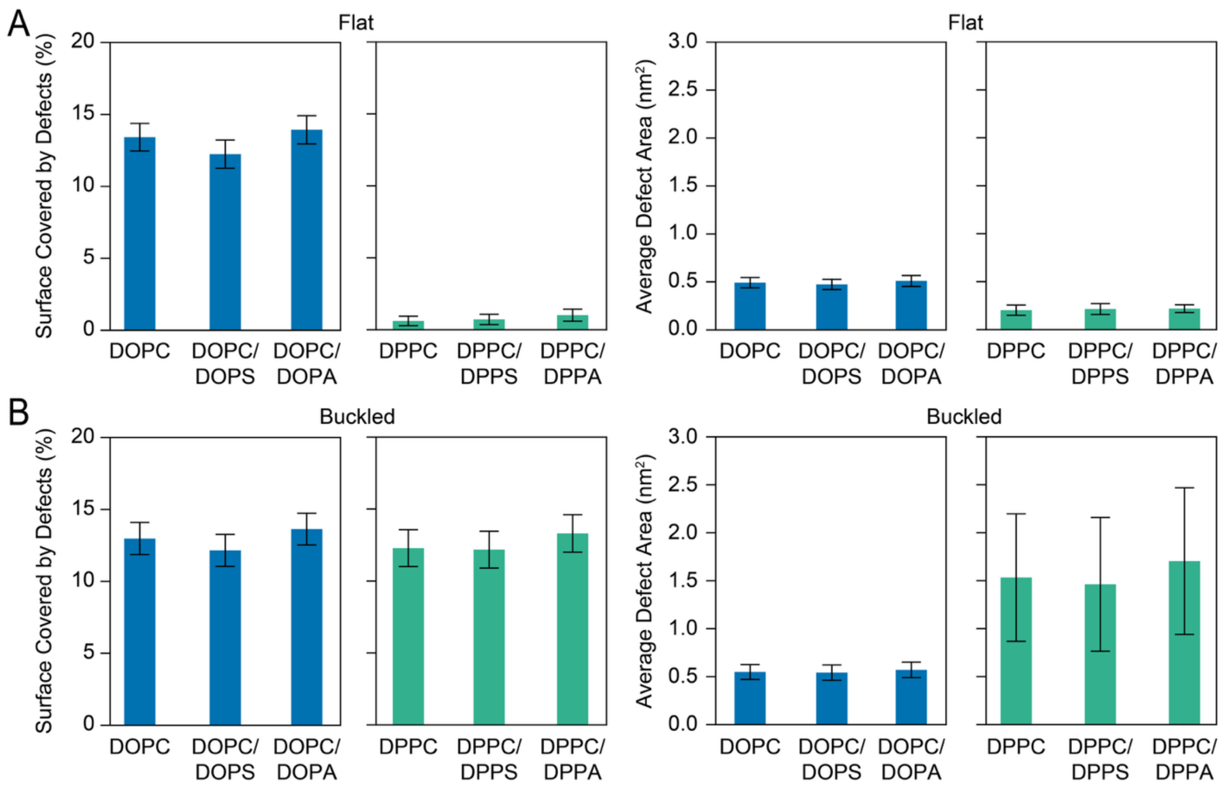


**Figure S9.** (A) Defect coverage and average defect area across the entire membrane surface for flat configurations. (B) Defect coverage and average defect area across the entire membrane surface for buckled configurations. Error bars in (A) and (B) represent the standard deviation calculated over all frames of the simulation.

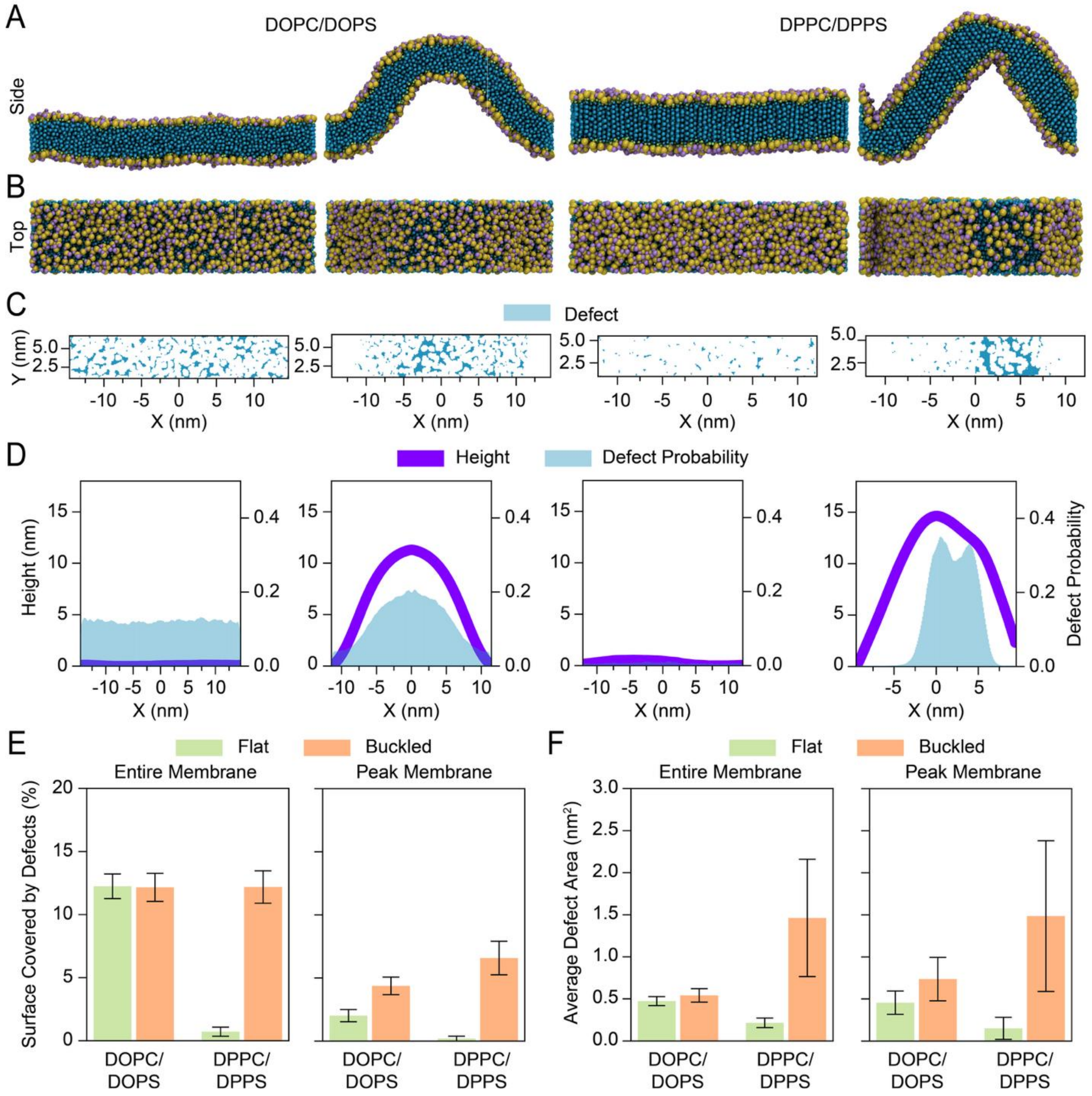


**Figure S10.** Representative (A) side and (B) top views of flat and mechanically buckled coarse-grained DOPC/DOPS (liquid-phase) and DPPC/DPPS (gel-phase) bilayers. (C) Lateral maps of defect locations for the representative bilayer above. (D) Spatial probability distribution of defects across the membrane. (E) Defect coverage and (F) average defect area across the membrane surface or restricted to the high-curvature crest region for flat and buckled configurations. Error bars in (E) and (F) represent the standard deviation calculated over all frames of the simulation.

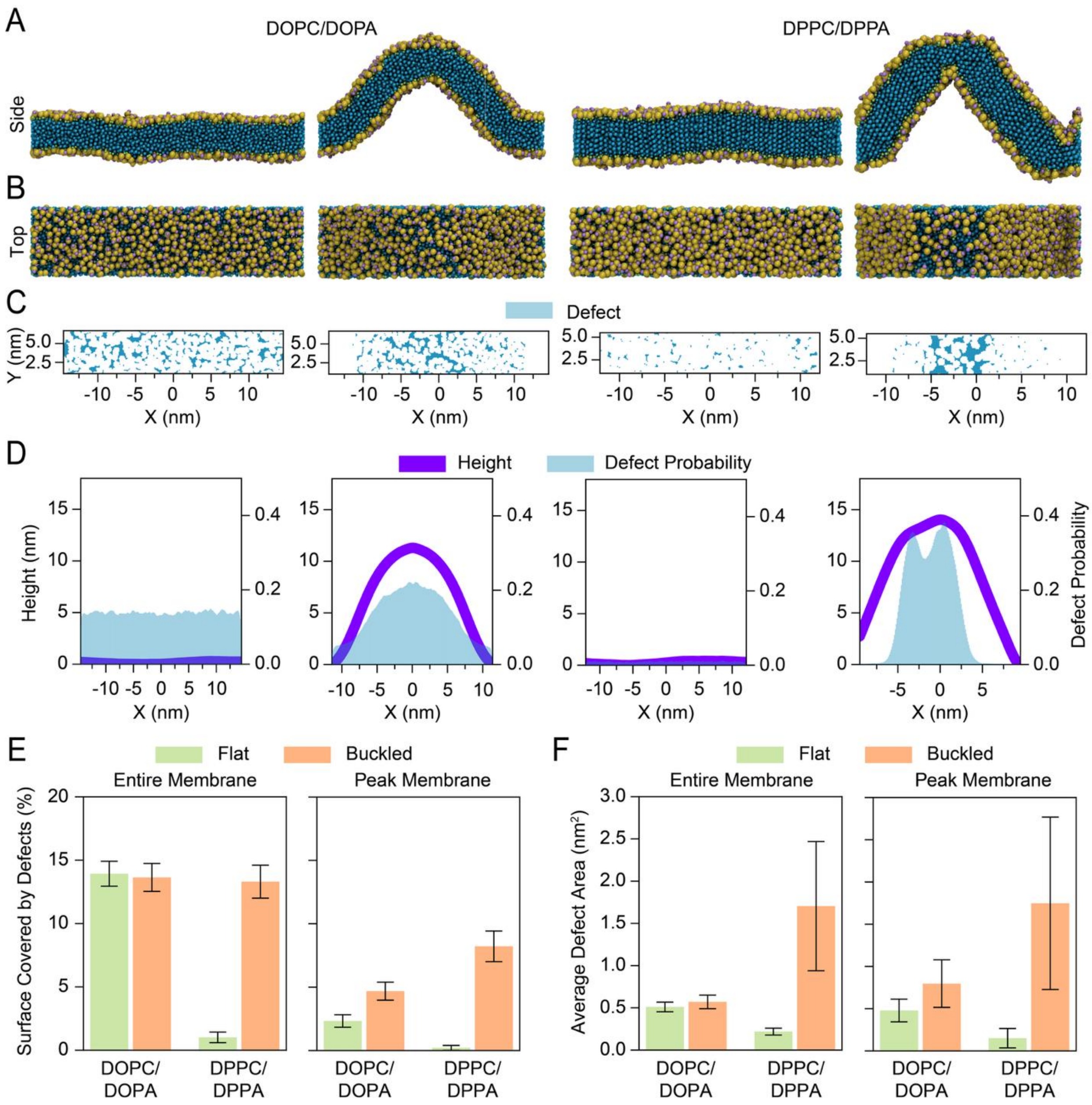


**Figure S11.** Representative (A) side and (B) top views of flat and mechanically buckled coarse-grained DOPC/DOPA (liquid-phase) and DPPC/DPPA (gel-phase) bilayers. (C) Lateral maps of defect locations for the representative bilayer above. (D) Spatial probability distribution of defects across the membrane. (E) Defect coverage and (F) average defect area across the membrane surface or restricted to the high-curvature crest region for flat and buckled configurations. Error bars in (E) and (F) represent the standard deviation calculated over all frames of the simulation.

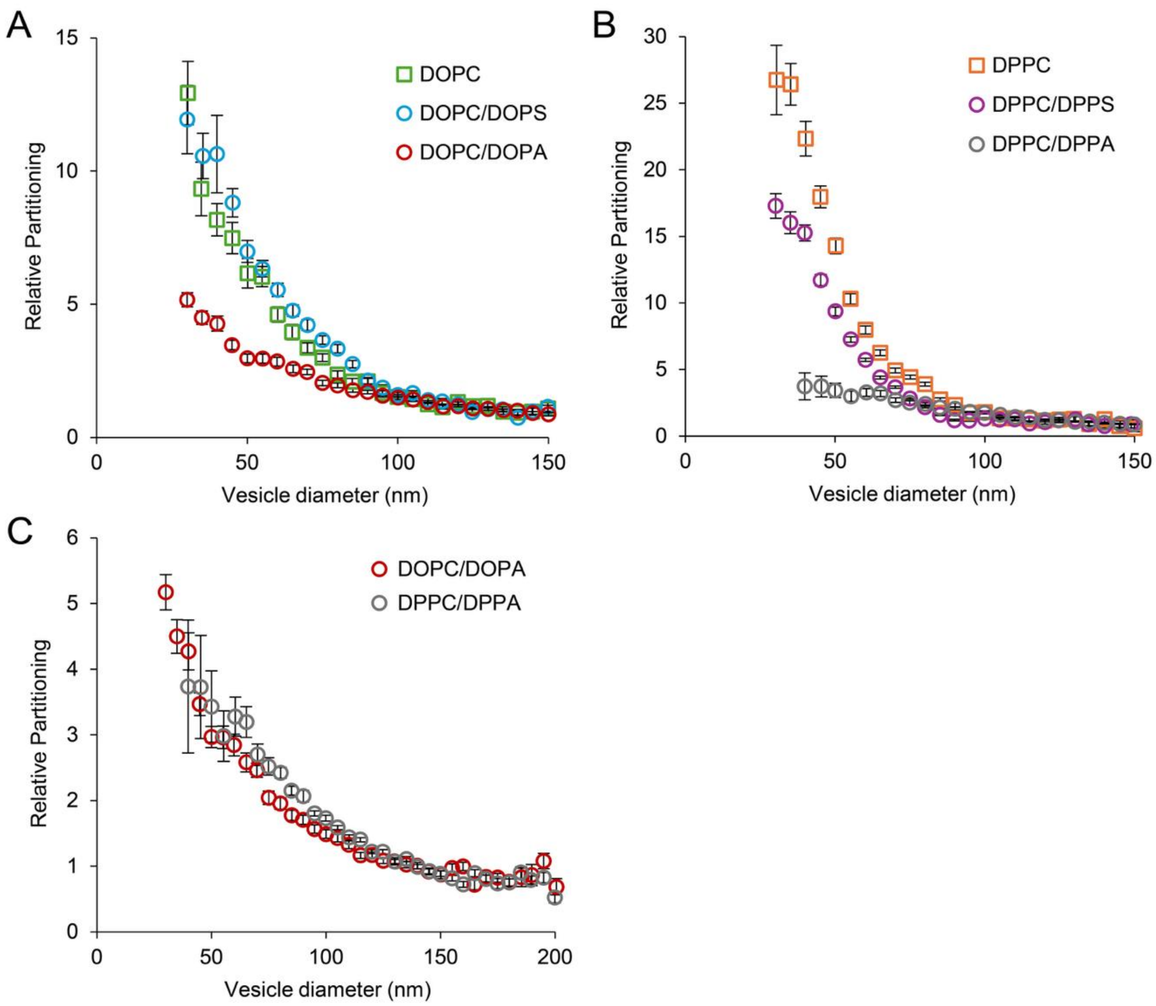


**Figure S12. Curvature sensitivity is diminished in liquid- and gel-phase vesicles with the incorporation of 25 mol% PA lipid.** (A-C) Relative partitioning of αSyn to vesicles as a function of vesicle diameter. Binding density was normalized to the average protein density on vesicles with diameters between 130-150 nm. Error bars represent the 99% confidence interval of the mean for all panels. PC/PS and PC/PA mixtures were combined at a molar ratio of 3:1 PC:anionic lipid. For the DPPC/DPPA data, vesicle populations in the 20 – 35 nm diameter range were overly sparse, thus unreliable for quantification. Therefore, data in this size range was omitted from the DPPC/DPPA plots in this figure.

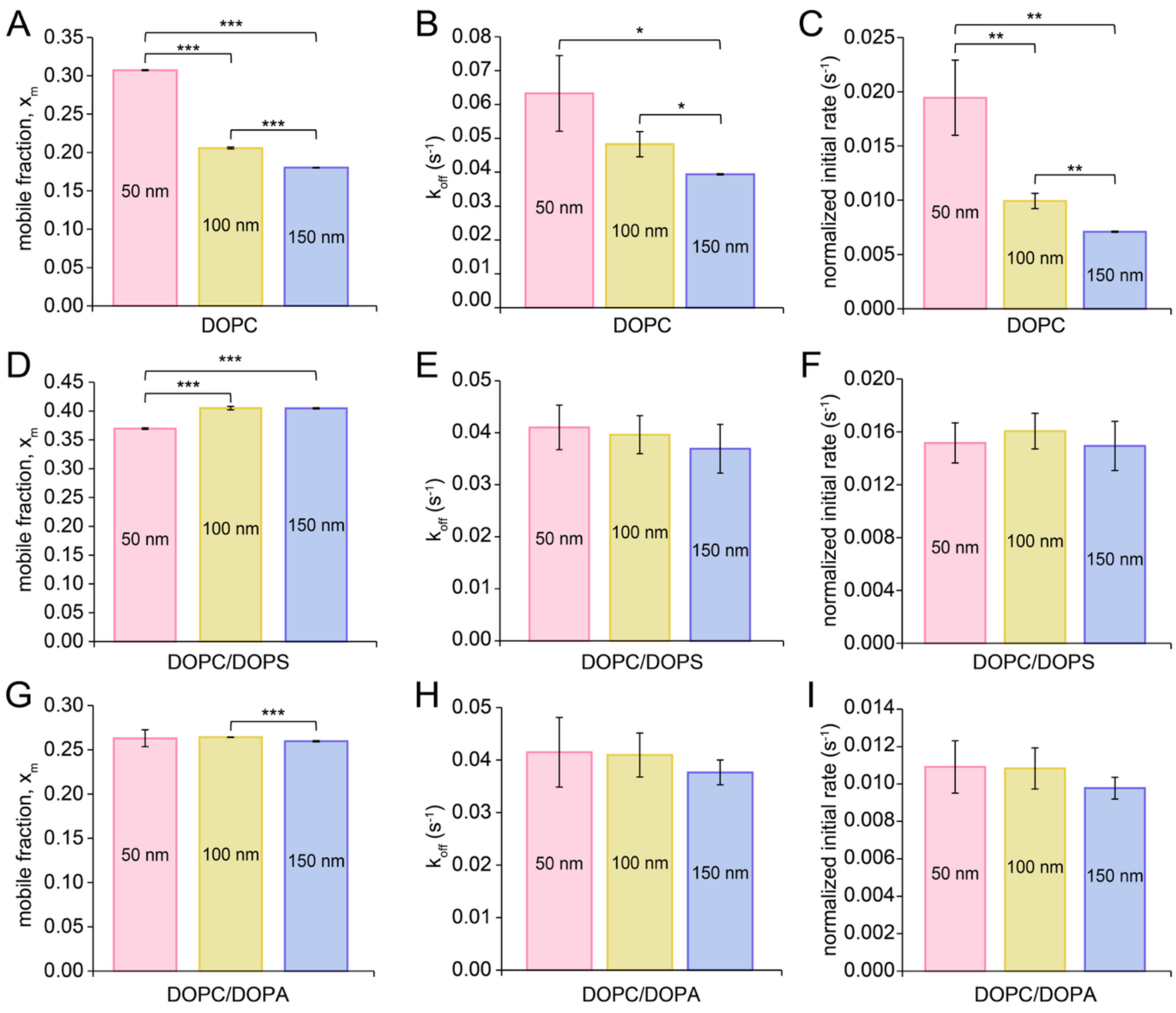


**Figure S13. Raw values for dynamic exchange kinetic parameters in liquid-phase membranes.** (A, D, G) Fraction of αSyn that is mobile on the membrane surface after photobleaching ($x_m$). (B, E, H) Dissociation rate constant ($k_{off}$) of αSyn from the membrane after photobleaching. (C, F, I) Normalized initial rate of αSyn dissociation, defined as the product of $x_m$ and $k_{off}$. Error bars represent the standard deviation (N = 3). * corresponds to $P < 0.05$, ** corresponds to $P < 0.01$, and *** corresponds to $P < 0.001$ as determined by unpaired Student's t-test.

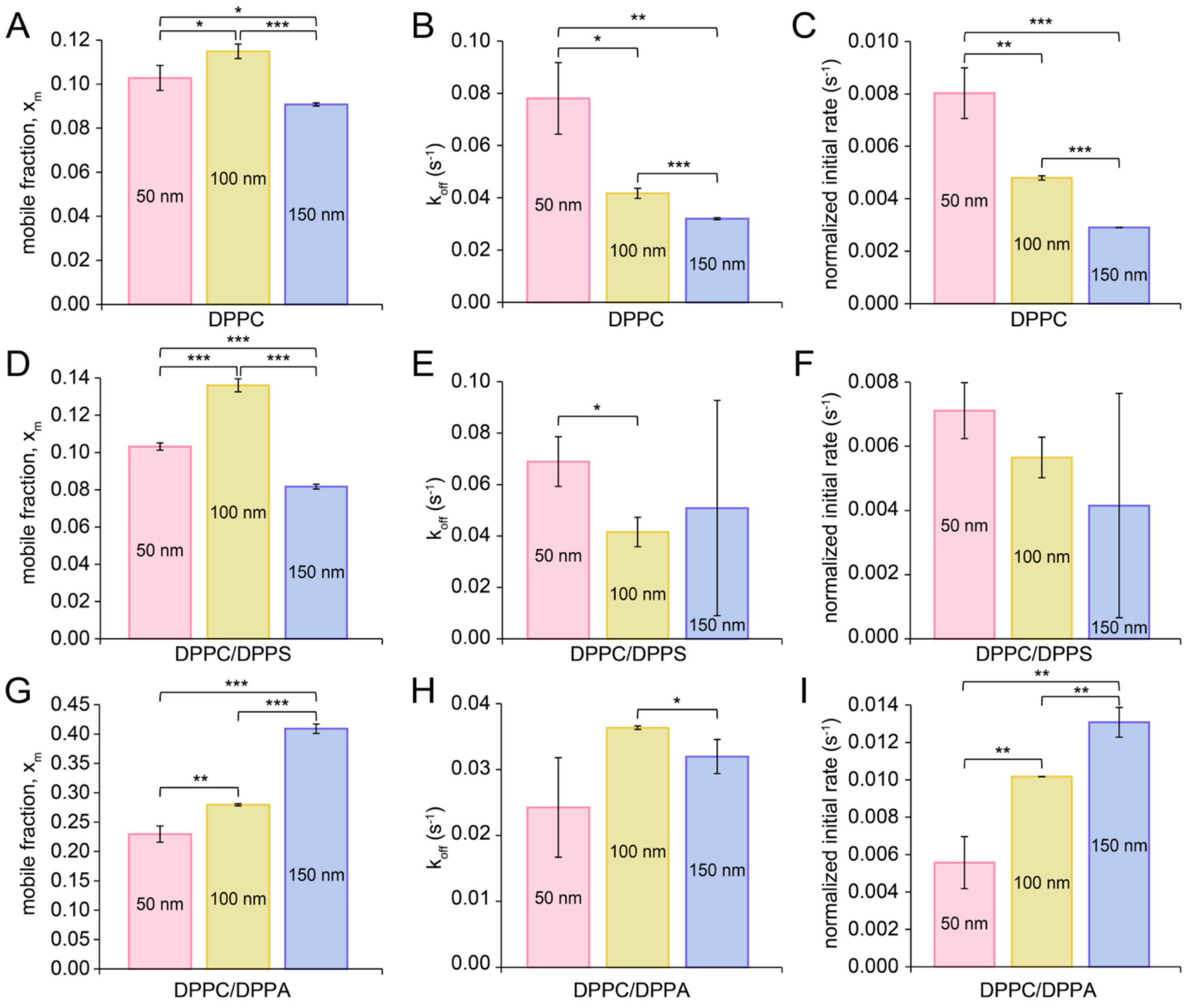


**Figure S14. Raw values for dynamic exchange kinetic parameters in gel-phase membranes.** (A, D, G) Fraction of αSyn that is mobile on the membrane surface after photobleaching ($x_m$). (B, E, H) Dissociation rate constant ($k_{off}$) of αSyn from the membrane after photobleaching. (C, F, I) Normalized initial rate of αSyn dissociation, defined as the product of $x_m$ and $k_{off}$. Error bars represent the standard deviation (N = 3). * corresponds to $P < 0.05$, ** corresponds to $P < 0.01$, and *** corresponds to $P < 0.001$ as determined by unpaired Student's t-test.

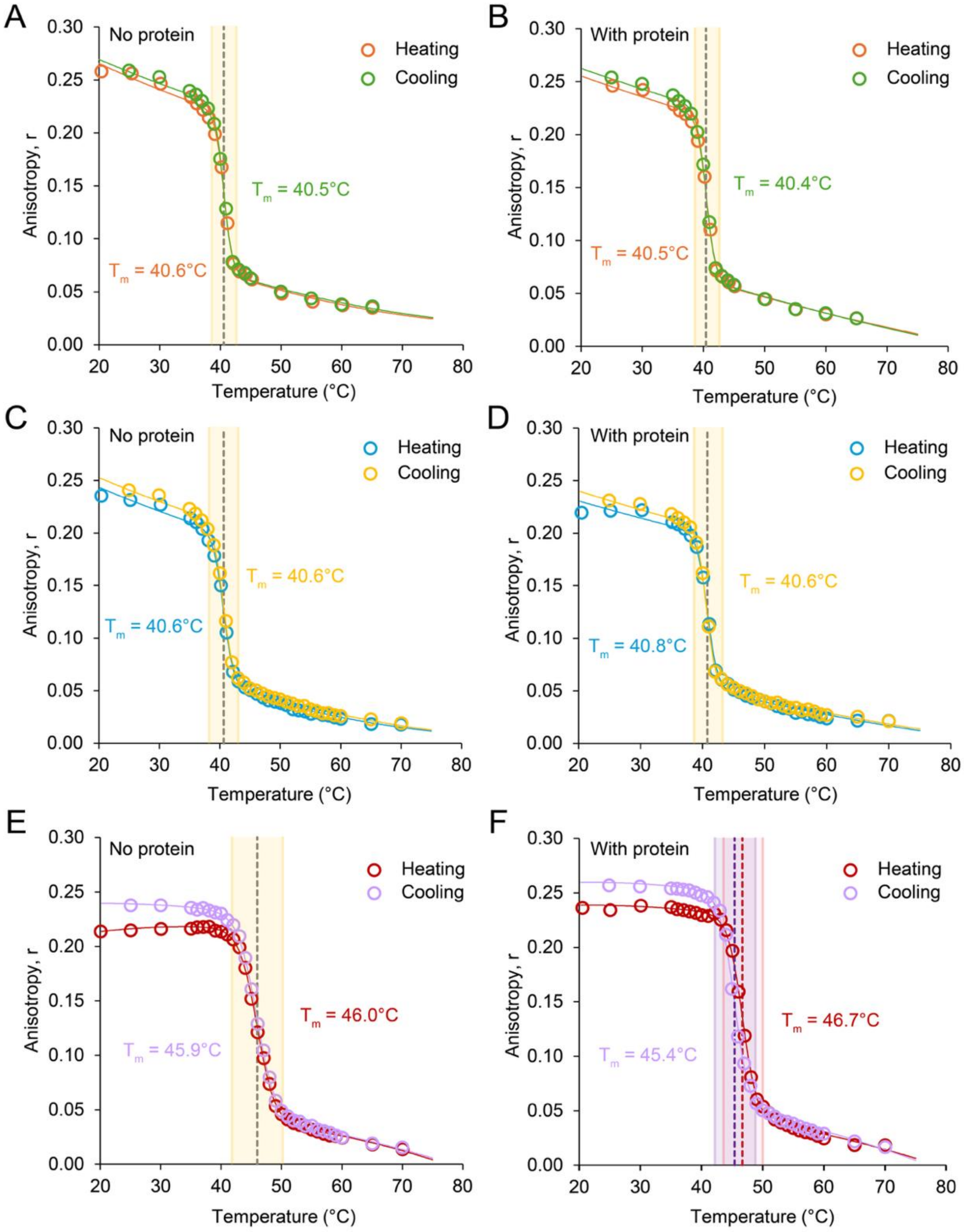


**Figure S15. Fluorescence anisotropy measurements capture lipid melting and unmelting through heating and cooling, respectively. The presence of αSyn does not substantially alter lipid phase behavior.** Fluorescence anisotropy curves for (A) DPPC vesicles, (B) DPPC vesicles with αSyn, (C) 3:1 DPPC/DPPS vesicles, (D) 3:1 DPPC/DPPS vesicles with αSyn, (E) 3:1 DPPC/DPPA vesicles, and (F) 3:1 DPPC/DPPA vesicles with αSyn. A 250:1 ratio of lipids to protein was used in experiments containing αSyn. Equation 6 was used to perform fits on the data. The dashed lines represent the $T_m$. The shaded region represents the boundary region for phase transition.

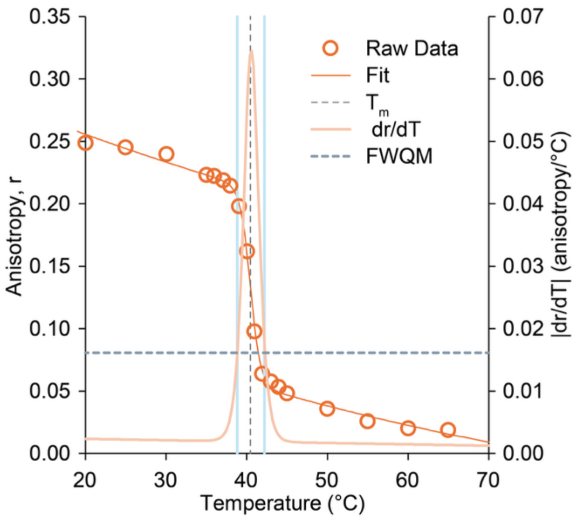


**Figure S16.** Representative anisotropy curve used to determine phase transition boundary regions. A Gaussian curve was obtained by taking the derivative of the anisotropy curve. The full-width quarter max values of the Gaussian curve were used to determine the upper and lower temperature boundaries (blue lines) for the lipid phase transitions.

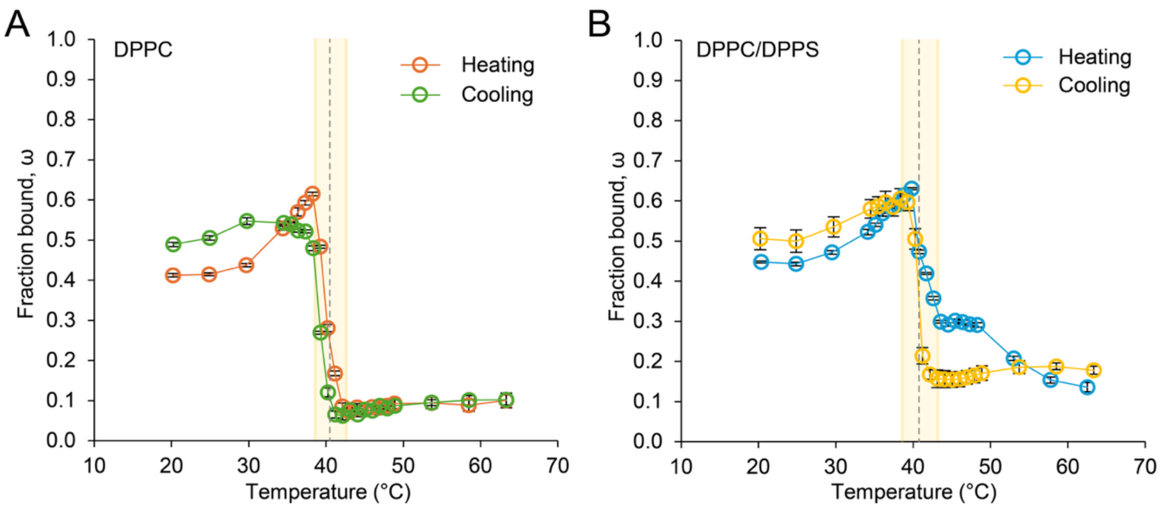


**Figure S17. Heating and cooling CD temperature ramp curves for DPPC and DPPC/DPPS membranes.** Binding curve for αSyn to (A) DPPC and (B) 3:1 DPPC/DPPS SUVs as the sample is heated and cooled. Data is obtained from CD spectroscopy. Error bars represent the SEM for the regression parameter ω at each temperature. The dashed line represents the $T_m$. The shaded regions represent the boundary regions for phase transition.

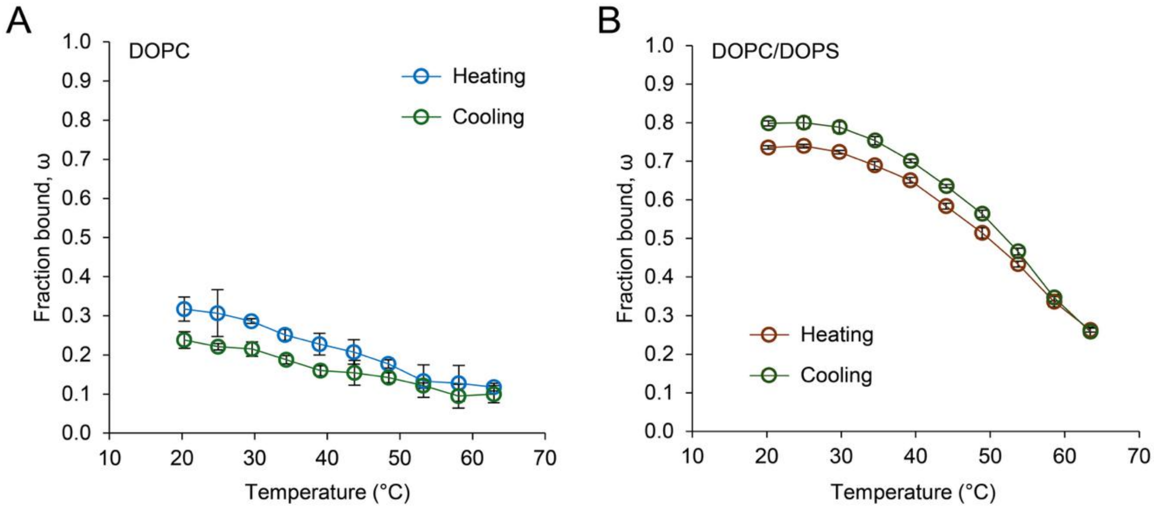


**Figure S18. Heating and cooling CD temperature ramp curves for DOPC and DOPC/DOPS membranes.** Binding curve for αSyn to (A) DOPC and (B) 3:1 DOPC/DOPS SUVs obtained from CD spectroscopy. Error bars represent the SEM for the regression parameter ω at each temperature.

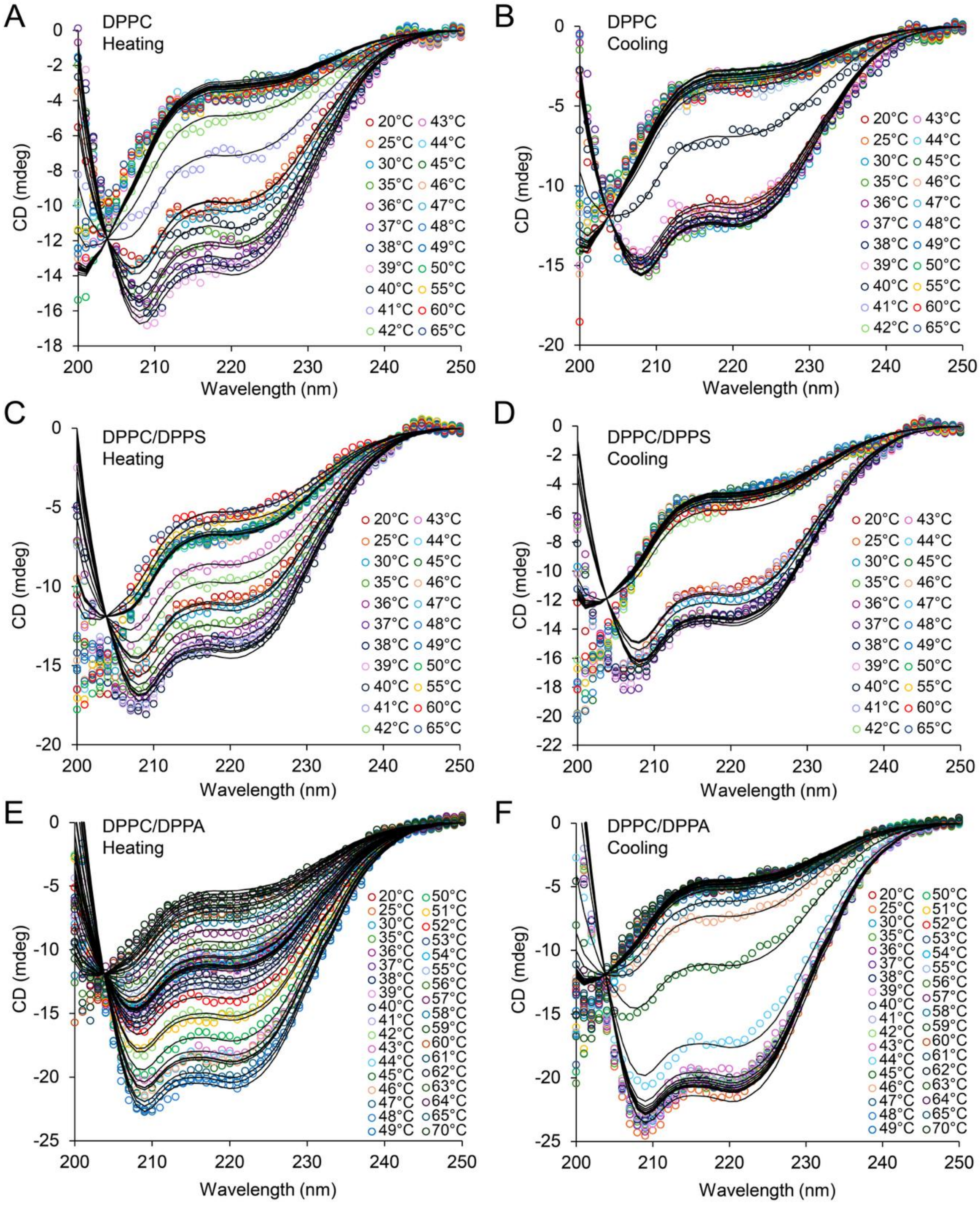


**Figure S19.** Circular dichroism spectra of αSyn bound to (A) DPPC vesicles as the sample is heated, (B) DPPC vesicles as the same sample is cooled, (C) 3:1 DPPC/DPPS vesicles as the sample is heated, (D) 3:1 DPPC/DPPS vesicles as the same sample is cooled, (E) 3:1 DPPC/DPPA vesicles as the sample is heated, and (F) 3:1 DPPC/DPPA vesicles as the same sample is cooled.

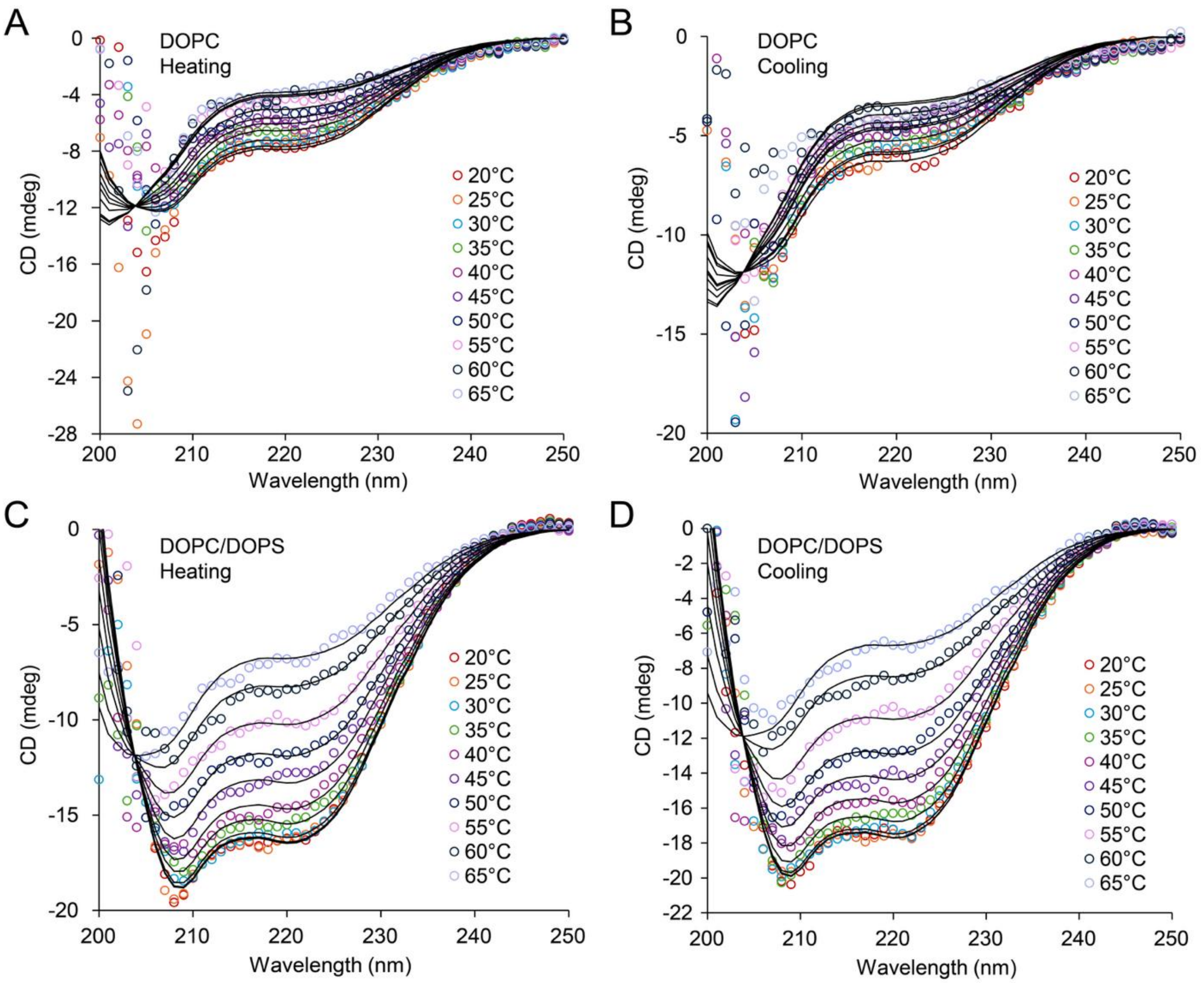


**Figure S20.** Circular dichroism spectra of αSyn bound to DOPC vesicles as the sample is (A) heated and (B) cooled, and αSyn binding to 3:1 DOPC/DOPS as sample is (C) heated and (D) cooled.

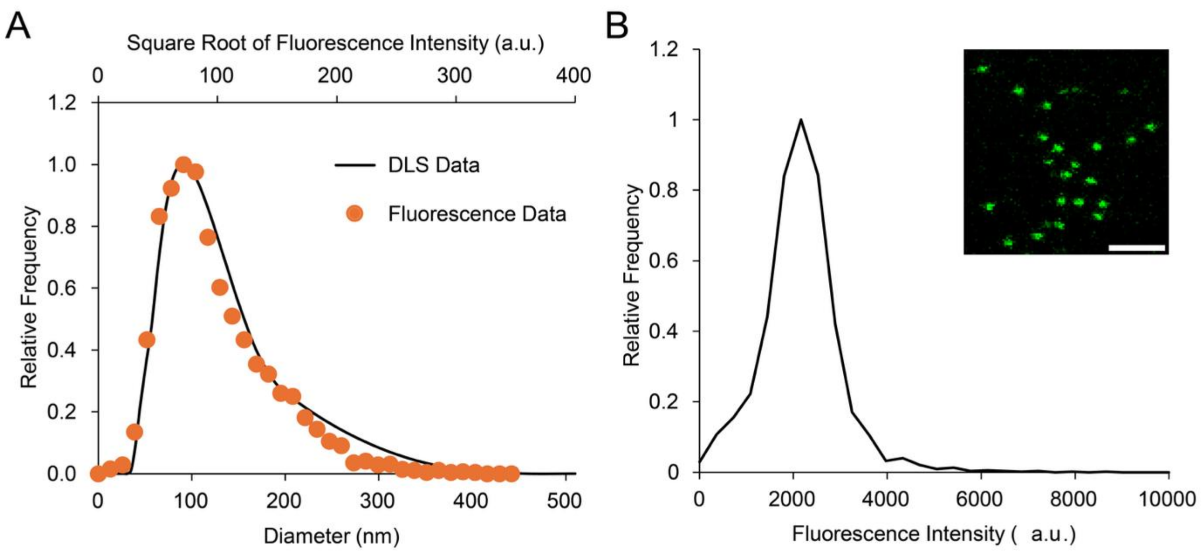


**Figure S21.** (A) Representative DLS/fluorescence intensity calibration plot for fluorescent SUVs. (B) Representative single-molecule fluorescence intensity distribution for fluorescent αSyn. Scale bar represents 2 μm.

**Table S1:** Composition for each system

| Lipids | System compositions (mol%) | PC lipids | PS lipids | PA lipids | Water | $Na^+$ | $Cl^-$ | Pressure strength (bar) | Strain |
|---|---|---|---|---|---|---|---|---|---|
| DOPC | 100 | 800 | 0 | 0 | 45988 | 580 | 580 | 3 | .199 |
| DPPC | 100 | 800 | 0 | 0 | 46032 | 580 | 580 | 35 | .199 |
| DOPC:DOPS | 75:25 | 600 | 200 | 0 | 45788 | 780 | 580 | 3 | .200 |
| DPPC:DPPS | 75:25 | 600 | 200 | 0 | 45836 | 780 | 580 | 38 | .199 |
| DOPC:DOPA | 75:25 | 600 | 0 | 200 | 46168 | 780 | 580 | 3 | .199 |
| DPPC:DPPA | 75:25 | 600 | 0 | 200 | 46172 | 780 | 580 | 30 | .199 |

**Table S2:** Major software and versions used in this study

| Software | Version |
|---|---|
| Gromacs[1] | 2025.2 |
| MDAnalysis[2; 3] | 2.10.0 |
| numpy[4] | 2.4.0 |
| Scipy[5] | 1.16.3 |
| matplotlib[6] | 3.10.8 |
| insane[7] | 1.2.0 |
| pandas[8; 9] | N/A |
| VMD[10] | 1.9.4a55 |
| MemDMA[11] | 1.0 |
| Asyn-phase-simulations[12] | N/A |

**References**


1. Gromacs 2025.2 manual. 2025. [accessed 2026]. https://zenodo.org/records/15387070.
2. Gowers RJ, Linke M, Barnoud J, Reddy TJE, Melo MN, Seyler SL, Domański J, Dotson DL, Buchoux S, Kenney IM et al. 2016. Mdanalysis: A python package for the rapid analysis of molecular dynamics simulations. SciPy.
3. Michaud-Agrawal N, Denning EJ, Woolf TB, Beckstein O. 2011. Mdanalysis: A toolkit for the analysis of molecular dynamics simulations. Journal of Computational Chemistry. 32(10):2319-2327.
4. Harris CR, Millman KJ, Van Der Walt SJ, Gommers R, Virtanen P, Cournapeau D, Wieser E, Taylor J, Berg S, Smith NJ et al. 2020. Array programming with numpy. Nature. 585(7825):357-362.
5. Virtanen P, Gommers R, Oliphant TE, Haberland M, Reddy T, Cournapeau D, Burovski E, Peterson P, Weckesser W, Bright J et al. 2020. Scipy 1.0: Fundamental algorithms for scientific computing in python. Nature Methods. 17(3):261-272.
6. Matplotlib: Visualization with python. 2025. v3.10.8. Zenodo; [accessed 2026].
7. Wassenaar TA, Ingólfsson HI, Böckmann RA, Tieleman DP, Marrink SJ. 2015. Computational lipidomics with insane: A versatile tool for generating custom membranes for molecular simulations. Journal of Chemical Theory and Computation. 11(5).
8. Pandas-dev/pandas: Pandas. 2025. v3.0.0cr1. Zenodo; [accessed 2026].
9. McKinney W. 2010. Data structures for statistical computing in python. SciPy.
10. Humphrey W, Dalke A, Schulten K. 1996. Vmd: Visual molecular dynamics. Journal of Molecular Graphics. 14(1).
11. Ctleelab/memdma: Memdma v1.0. 2026. Zenodo; [accessed 2026]. https://zenodo.org/records/20076551.
12. Buckled and planar membrane phase data. 2026. Zenodo; [accessed 2026]. https://zenodo.org/records/20073648.
